# PCP Auto Count: A Novel Fiji/ImageJ plug-in for automated quantification of planar cell polarity and cell counting

**DOI:** 10.1101/2024.01.30.578047

**Authors:** Kendra L. Stansak, Luke D. Baum, Sumana Ghosh, Punam Thapa, Vineel Vanga, Bradley J. Walters

**Author notes:** Corresponding author (BW).

## Abstract

**Background:** During development, planes of cells give rise to complex tissues and organs. The proper functioning of these tissues is critically dependent on proper inter- and intra-cellular spatial orientation, a feature known as planar cell polarity (PCP). To study the genetic and environmental factors affecting planar cell polarity investigators must often manually measure cell orientations, which is a time-consuming endeavor.

**Methodology:** To automate cell counting and planar cell polarity data collection we developed a Fiji/ImageJ plug-in called PCP Auto Count (PCPA). PCPA analyzes binary images and identifies “chunks” of white pixels that contain “caves” of infiltrated black pixels. Inner ear sensory epithelia including cochleae (P4) and utricles (E17.5) from mice were immunostained for βII-spectrin and imaged on a confocal microscope. Images were preprocessed using existing Fiji functionality to enhance contrast, make binary, and reduce noise. An investigator rated PCPA cochlear angle measurements for accuracy using a 1-5 agreement scale. For utricle samples, we directly compared PCPA derived measurements against manually derived angle measurements using concordance correlation coefficients (CCC) and Bland-Altman limits of agreement. Finally, PCPA was tested against a variety of images copied from publications examining PCP in various tissues and across various species.

**Results:** PCPA was able to recognize and count 99.81% of cochlear hair cells (n = 1,1541 hair cells) in a sample set, and was able to obtain ideally accurate planar cell polarity measurements for over 96% of hair cells. When allowing for a <10° deviation from “perfect” measurements, PCPA’s accuracy increased to >98%. When manual angle measurements for E17.5 utricles were compared, PCPA’s measurements fell within −9 to +10 degrees of manually obtained mean angle measures with a CCC of 0.999. Qualitative examination of example images of Drosophila ommatidia, mouse ependymal cells, and mouse radial progenitors revealed a high level of accuracy for PCPA across a variety of stains, tissue types, and species. Altogether, the data suggest that the PCPA plug-in suite is a robust and accurate tool for the automated collection of cell counts and PCP angle measurements.

## Introduction

The proper orientation of cells with regard to anatomical axes, neighboring cell orientation, and overall system organization is of critical importance for proper development and function in nearly all multicellular organisms. During metazoan development, immature sheets of cells respond to polarization cues that provide the organizational direction necessary for the formation of complex tissues, organs, and systems. The study of coordinated cell polarization in these two dimensional sheets— known as planar cell polarity (PCP)—has greatly benefitted from the improvement of molecular and gene targeting methodology developed over the past few decades, and there has been considerable scientific interest in studying PCP pathways in a variety of developing tissues and model organisms [1–6]. Indeed, several factors that regulate PCP have been identified, including several “core” PCP proteins that appear to be largely conserved [7,8]. In addition, several recent publications show that there are novel PCP factors to be discovered [9,10], some that may be specific to certain tissue types or species [11,12], and there is also a growing list of functions for many PCP proteins beyond mere polarization [13,14]. As this area of study continues to grow, new tool development and adoption of streamlined methodologies will be highly beneficial.

A major technical limitation in the study of PCP is that the manual characterization and collection of cellular polarity data is incredibly labor intensive. Most types of tissue contain thousands-to-millions of cells, making it nearly impossible to conduct analyses on the entire cell population within an organ. Many investigators elect to quantify cell orientations from a limited number of sampling areas or cells and then extrapolate broader regional conclusions based on this data. Even with such random and limited sampling, statistical power and experimental rigor often require the measurement of hundreds of cells, which is still intensive and vastly time consuming for investigators taking manual measurements. Furthermore, limited sampling techniques may lead investigators to draw incorrect regional assumptions, particularly in cases where cell type ratios or cell density varies based on anatomical location [15] or developmental age [16]. Beyond the extensive time commitment necessary to collect PCP data, manual quantification of planar cell polarity—even by blinded investigators— is subject to human error and potential bias. Due to the limitations inherent in manual cell quantification, there has been a growing interest in the development of automated or semi-automated processes that would allow researchers to collect PCP data quickly and accurately. Automation of PCP data collection should provide tremendous savings in terms of time and human resources, with the added benefit of minimizing variability and bias.

To address the need for a reliable and user-friendly approach to automating PCP data collection, we have developed a user-friendly plug-in tool suite called PCP Auto Count (PCPA). PCPA automates cell quantification and collects PCP data in the form of angle measurements from two-dimensional micrographs. PCPA is integrated with the widely used open-source software FIJI (Fiji Is Just ImageJ) [17] and allows researchers to customize data collection parameters through a simple graphical user interface. To test the efficacy of PCPA we utilized a number of confocal fluorescent micrographs taken from murine inner ear sensory epithelia and compared PCP data collected with PCPA against PCP data collected through traditional manual quantification methods.

Inner ear auditory and vestibular sensory epithelia have become prominent models for the investigation of how disruption to developmental factors can affect PCP [9,18,19]. The main sensory cell type of the inner ear—known as hair cells—are activated when stereocilia protruding from the apical surface of the cell are pushed by mechanical forces in the direction of the cell’s primary cilium or kinocilium. Mechano-electric transduction channels are opened, depolarizing the cell and triggering neurotransmitter release onto the vestibulocochlear nerve. In order to be maximally sensitive to deflecting forces, sensory hair cells develop specific spatial orientations during late embryonic and early postnatal ages. Disruption to developmental factors influencing PCP have been shown to lead to deficits in hearing and balance [20–22]. Thus, we have developed the PCPA plug-in suite to automate the collection of cell polarity measurements and validated its utility extensively in cochlear and vestibular sensory epithelia. In addition, we demonstrate the ability of PCPA to obtain orientation measurements from published images of *Drosophila* ommatidia, murine ependymal cells, and murine radial glia. Together, the data suggest that PCPA is a reliable and accurate plug-in suite that performs at least as well as manual quantification, and has significant potential to streamline and expand planar cell polarity analyses in multiple cell types and tissue models.

Finally, though the PCPA plug-in was primarily developed to calculate cell polarity measurements, it also provides broader experimental applicability through its ability to quantify cell numbers. Across many areas of study in the biological sciences, the counting of cells is necessary to understand the effects of experimental manipulations. Investigators often wish to quantify changes in RNA or protein expression *in situ*, or on the processes of cell survival, proliferation, and differentiation by counting cells expressing certain markers. In the inner ear, sensory cells can be damaged or lost due to environmental factors, genetic predisposition, and the normal aging process. Many research questions related to sensory cell protection or regenerative strategies use cell population counts as an important outcome metric, and this approach [23–25] is subject to the same limiting factors described above (time investment and potential bias). Though not heavily emphasized here, our results demonstrate that PCPA can be used to quantify cell numbers in addition to angles of orientation, or independently for samples where cell polarity information is not needed.

## Results

### Optimization of image acquisition: high signal-to-noise promotes reliable conversion to black and white

PCPA was designed to analyze single plane, two-dimensional images that are binary (black and white) with the cells represented by white pixels and a concave or enclosed directional point of interest comprised of black pixels. In practice, micrographs are not generally acquired in this manner, often being acquired in color or grayscale and in some cases as confocal z-stacks. While polychromatic and grayscale images are useful for quantification methods relying on human decision making, automated algorithmic quantification is easier to accomplish and requires less processing power when the image information is limited to binary pixel composition. PCPA was therefore designed for analysis of binary images which reduces computation and improves speed of execution. As such, multicolor or grayscale images need to be converted to binary images which can be easily accomplished using existing Fiji features. Similarly, z-stacks can be readily cropped or projected onto a two-dimensional image. Over many iterations we have established that reliable image acquisition and a limited number of preprocessing steps can decrease background noise and yield the best flattening and binary conversion from a grayscale or color image.

Our image acquisition parameters were established using murine cochlear (P4) hair cells immunolabeled for the commonly used sensory hair cell marker βII-spectrin. In order to capture images that required minimal preprocessing to create quality binary images, the βII-spectrin channel was imaged with the detector gain set to approach or slightly exceed saturation for immunopositive pixels. We have found that binary images with the best fidelity to detail are created from brightly imaged micrographs obtained at relatively high resolution (minimum 1024 x 1024 pixels field of view), at high magnification (40X to 60X microscope objective), and using a robust antibody with high signal-to-background fluorescence ratio. In P4 cochlear samples, βII-spectrin signal intensity appeared brighter in inner hair cells compared to outer hair cells. Uneven signal intensities such as these sometimes led to issues when thresholding images to binary. Thus, relatively even levels of fluorescence across all cells in a field should be attained if possible. Where such evenness in intensity was not achievable in images from P4 cochleae, inner and outer hair cells were separated into different images and subsequently preprocessed and analyzed with great success.

While it is recommended to maximize signal-to-noise ratio (SNR) of fluorescence intensities, and to oversaturate as necessary to ensure homogeneity across cells for ease of use of PCPA, we also tested PCPA using images that were collected for prior experiments using standard imaging parameters where pixel intensities were not maximized or normalized, thus demonstrating how PCPA could be useful in situations where researchers wish to analyze existing micrographs or to maintain existing imaging parameters for other reasons. We utilized micrographs of murine vestibular (E17.5) hair cells, from our laboratory, that were immunolabeled for βII-spectrin and imaged prior to the development of PCPA. We also tested PCPA on images from other laboratories in their previously published research. These images did not meet our resolution requirements (< 1024 x 1024 pixels) so we artificially increased the resolution in Fiji using “Image > Adjust > Size…” and set the size to meet or exceed 1024 pixels on the shortest edge of the image.

### Inbuilt Fiji functionality can be used to pre-process micrographs for PCPA analysis

For P4 cochlear images obtained using optimal microscopy parameters described above, we found that further pre-processing was often not necessary and we could move straight to creating binary images from these micrographs. For any images that did not meet the criteria defined above (high SNR and flat 2D image), we found that pre-processing steps completed using existing features in Fiji enhanced the quality of subsequently created binary images, and ultimately resulted in nearly all positively labeled cells from the micrographs to be processed successfully by PCPA. We briefly describe the preprocessing steps used below, and a detailed explanation of pre-processing settings can be found in the PCPA user manual which can be obtained within the PCPA plug-in (or at https://github.com/WaltersLabUMC/PCP-Auto-Count.git).

First, to create 2D images from 3D (z-stack) micrographs, the Fiji function “Image Stacks > Z project…” allowed for projection using a variety of methods. Of these, the maximum intensity projection worked best for most images, though the median intensity function sometimes yielded better results. In cases where SNR was low, PCPA performance was improved by selecting “Process > Subtract background…”. Standard Fiji functions were not always sufficient to correct high heterogeneity of signal within particularly poor images. In these cases, we corrected image intensity using the BioVoxxel plug-in tool suite (www.biovoxxel.de). Specifically, the “pseudo flat-field correction” option and/or “convoluted background subtraction” option improved homogeneity of signal intensities and allowed further background subtraction if needed after flat-field correction. After background subtraction (if applicable), images were converted to binary, most commonly by using the Fiji function “Image > Adjust > Threshold…,” where users manually set the threshold point. Alternatively, for images that were of sufficient quality and homogeneity, Fiji’s automated threshold options could be used (e.g. “Process > Binary > Make Binary”).

When making an image binary, nonspecific staining could be intense enough to be included as white pixels. While distinct non-specific white pixels can be size excluded (see below), white pixels that arise in direct contact with cells of interest could lead to erroneous calculations of the centers of the cells during PCPA analysis.

Furthermore, it is generally beneficial to PCPA processing to eliminate spurious concavities from cells, i.e. any indentations or holes that are not the feature to be measured for directionality. While allowing such features to remain would likely only slow PCPA analysis by a few seconds, it could lead to occasionally erroneous angle measurements. Thus, it is highly recommended to use the Fiji option “Process > Noise > Remove outliers…” to remove these non-specific features. For the images used in this report, noise removal between 6-12 pixels was generally sufficient.

### PCPA development and algorithm workflow

#### PCPA algorithm overview, download and source code

The feature of interest in most PCP studies is often a cell, or surface of a cell, but may alternatively be a cellular structure, aggregate of cells, aggregate of structures, or other features that can vary by tissue type. As such, we have elected to use generalized descriptive terms in the development and functioning of PCPA, which should be broadly applicable. The primary features of interest, usually large planar surfaces (often the apical surfaces of cells) are termed “chunks”, while prominent inclusions indicative of direction are defined as “caves.” These chunk and cave configurations are present in many types of samples used for PCP research (e.g. v- or u-shaped morphology such as the arrangement of photoreceptors in *Drosophila* ommatidia, filiform papillae of the vertebrate tongue, actin-rich *Drosophila* wing hairs, or stereocilia bundles of mammalian outer hair cells). This organization of features also often arises from combinations of various features, such as the co-labeling of cell surfaces or borders and primary cilia (e.g. radial glia, ependymal cells, and inner ear hair cells). In many remaining cases, chunks and caves can be overlaid onto features of interest by investigators who circle or otherwise draw on top of an image to highlight the points of interest. The primary principle in the operation of PCPA is that the angles of orientation of cells or groups of cells in planar tissues can be measured using two reference points in each cell or cell aggregate; namely, the center of mass of the chunk and the center of mass of the cave. As a proof of principle, here we utilize cochlear and vestibular sensory hair cells where the apical surface (chunk) is immunolabeled with βII-spectrin and the commonly used orientation marker of hair cells, the fonticulus, is an unlabeled circular inclusion (cave). We also provide examples of PCPA’s utility on other cell types (e.g. *Drosophila* ommatidia, murine ependymal cells, and murine radial glia) taken from published PCP literature. Here, we provide a brief description of PCPA’s workflow and algorithmic processes, presented in operational order (Fig 1). The PCPA Fiji plug-in can be readily installed into Fiji from: https://sites.imagej.net/PCP-Auto-Count/ and detailed instructions for installing Fiji plug-ins can be found in the PCPA user manual or through Fiji’s wiki at https://imagej.net/plug-ins/updater or https://imagej.net/plug-ins/. A user manual with detailed explanations of the PCPA algorithm and function, along with PCPA’s open source code, can be found at https://github.com/WaltersLabUMC/PCP-Auto-Count.git.

**Fig 1.**
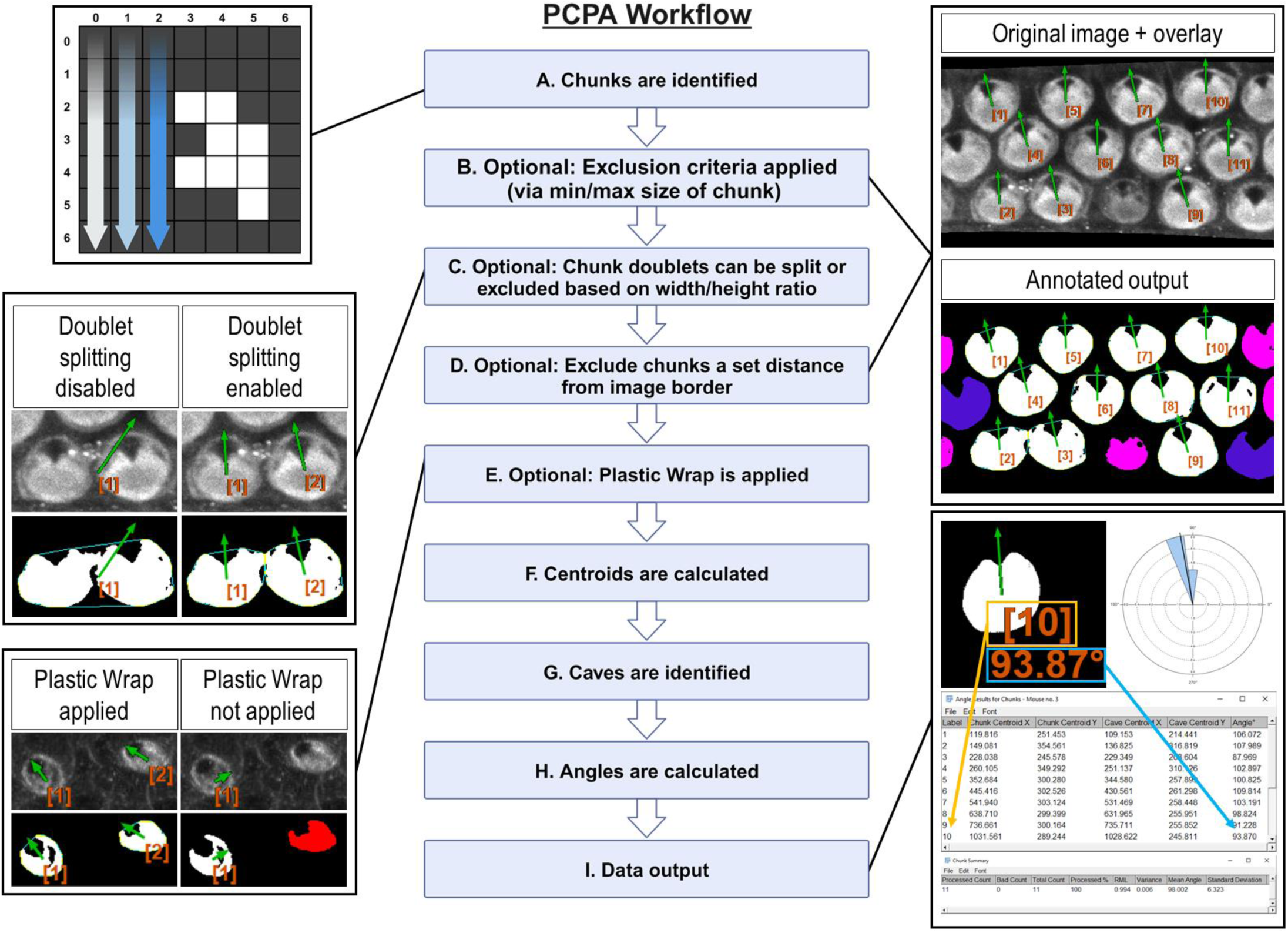
Overview of PCPA’s algorithmic flow (Created with BioRender.com) **(A)** PCPA identifies aggregates of white pixels in a binary image, termed chunks. **(B)** Chunks can be discarded from subsequent operations via an optional pixel size minimum and/or maximum (cells pseudo-colored pink). **(C)** PCPA has a limited capacity (maximum of two conjoined cells) to separate chunks that abut one another or overlap. Larger aggregates of chunks can be excluded by overall size or dimensions. **(D)** Cells that fall partially outside of the imaging frame can be discarded by setting an exclusion zone of a set number of pixels away from the edge of the image frame (cells pseudo-colored blue). **(E)** In the absence of Plastic Wrap, fonticuli that are not completely enclosed in βII-spectrin staining cannot be recognized by PCPA. Applying the “Plastic Wrap” option will add pixels to bridge the gap in the cell’s perimeter where the unenclosed fonticulus would be. **(F)** Each chunk’s center of mass is calculated and stored as (x, y) coordinates. **(G)** Inclusions of black pixels within chunks, termed caves, are identified and the cave best meeting selection criteria is identified as the directional cave of interest and the center of mass of the enclosed black pixels is calculated as (x, y) coordinates. **(H)** The angle of orientation for each chunk and its cave is calculated as the inverse tangent of the difference in (y) over the difference in (x). **(I)** PCPA outputs include annotated images specifying chunks excluded due to violating chunk size limits (cells pseudo-colored pink) or touching the border of the image (cells pseudo-colored blue). Cells are labeled in the annotated output and correspond with the angle results table, which provides a detailed output of (x, y) coordinates and angles for each chunk-cave pair that was quantified. Rose diagrams and summary statistics tables are also produced.

#### PCPA identifies white pixels in a binary image and aggregates adjoining white pixels into discrete chunks

PCPA will first identify all potential chunks in a binary image. To do so, the Cartesian (x, y) coordinates of all white pixels in the image are recorded. PCPA then systematically evaluates every white pixel to determine if it abuts any other white pixels. All abutting pixels are assigned as an aggregate representing an individual chunk (Fig 1A). The coordinates of all chunks are then stored for later use in calculating the center of mass of each chunk. Users can set optional upper and/or lower size limits to exclude chunks from analysis (Fig A in S1). This allows users to filter out spurious signal and to select for cells in a size range, which can be useful for filtering out immature cells (Fig 1B).

As the fusion of neighboring cells in micrographs is a common problem in automated image analysis [26–28], we developed an optional Doublet Splitting mode, where PCPA will separate two cells that touch or overlap by splitting the doublet mass in half, then treating each half as a discreet cell during subsequent steps (Fig B in S1). This optional function cannot recurse beyond separating two conjoined cells. and works most effectively when fused cells are somewhat reasonably aligned with the x or y axis of an image. In this way, users can avoid having to manually separate all conjoined cells in an image. Alternately, users can choose to discard all doublets (triplets, or larger aggregates) from further analyses via size exclusion if preferred (Fig 1C). After any optional doublet splitting or exclusion is completed, PCPA will then optionally exclude cells a set distance from the image border (Fig A in S1). Setting a border exclusion zone is recommended as this allows users to filter out cells that did not completely enter the imaging frame. After discarding all chunks meeting the user selected criteria, PCPA calculates the center of mass for each chunk. The chunk center of mass is then used as the point from which the vertex is created for subsequent PCP angle measurements (Fig 1F).

#### PCPA identifies inclusions of black pixels in chunks and subsequently assigns one cave as the directional point of interest

To identify candidates for the cave that will ultimately serve as the directional marker for PCP angle calculations, PCPA will first compile the coordinates of all caves in each chunk. A cave is defined as any black pixel, or aggregate of black pixels, fully contained within the body of a chunk (i.e. completely surrounded by white pixels constituting the chunk). If the primary PCP-indicating feature presents as unenclosed indentations (e.g. the u-shaped configuration of rhabdomeres in *Drosophila* ommatidia), we developed an optional feature called Plastic Wrap to enclose such concavities (Fig A and A’ in S2).

Plastic wrap identifies any breaks in the arc of a chunk’s perimeter due to indentations in the chunk, then subsequently adds a 1-pixel thick bridging line across these areas (Fig 1E). Algorithmically, Plastic Wrap runs immediately prior to chunk centroid calculations because the bridging pixels added by Plastic Wrap are considered part of the cell mass for subsequent chunk centroid calculations. For a more in-depth explanation of Plastic Wrap, see the PCPA user manual or source code. After all cave candidates of all chunks are identified (including caves created by Plastic Wrap), PCPA will designate one cave for each chunk that best meets user-defined selection criteria as the directional cave of interest. Users must set a selection criterion of either the largest cave in a chunk or the cave best meeting a directional criterion (i.e. northmost, etc.; Fig C & C’ in S2). This selection criteria can be combined with optional cave minimum and/or maximum size requirements, which will exclude any caves violating size requirements prior to cave of interest selection (Fig B in S2). PCPA will then calculate the center of mass for the cave of interest to use for subsequent PCP angle calculations (Fig 1G).

#### PCPA calculates angles and creates annotated data output and summary statistics

PCPA will calculate the inverse tangent of the chunk and cave centers of mass for each chunk/cave pair in the image (Fig 1H). Angle measurements are dependent on the scale and direction of the directional (XY) axis. Angle measurements can be calculated on a −180°/180° or 0°/360° axis where angles increase incrementally clockwise or counter-clockwise. Users are able to customize their axis settings in the PCPA options dialogue (Fig A-C in S3).Once angle measurements have been calculated, PCPA will output data tables and annotated images, explained as follows (Fig 1I). The Results Table consists of a data row for each chunk analyzed and contains: an identification number assigned to each chunk, the (x, y) coordinates of the chunk, and the angle measurement of the chunk-to-cave vector. The Chunk Summary table contains summary statistics for the data set, including: Processed Count (the number of cells where an angle was measured), Bad Count (the number of cells for which no angle was measured), Total Count (sum of Processed Count and Bad Count), Processed %, mean angle, resultant mean length (RML), circular variance, and circular standard deviation. The output also includes a circular data histogram known as a rose or windmill diagram. The Successfully Processed Chunks output image features annotations superimposed onto the binary image used for analysis, and includes each cell’s identification number, angle measurement, and a directional arrow derived from the angle measurement. This output also features pseudo-coloring to indicate if any chunks were excluded from analysis and why. Pseudocolor designations can be customized by the user so that each exclusion category is color specific (Fig E in S4). Lastly, an overlay image containing only the identification number, angle measurement, and directional arrow for all analyzed cells in the image is produced. This overlay image can be superimposed over top the original grayscale or color micrograph using inbuilt Fiji functionality (“Image > Overlay > Add image… Zero transparent”). Each of these outputs can be selected or deselected by users in the PCPA options dialog (Fig A-D in S4).

While the aforementioned functionality described above constitutes the main intended use of PCPA analysis using a single binary image, PCPA has two further functionalities of note. Firstly, PCPA can batch process multiple binary images, aggregate each image’s cell measurements into one data set, then produce a Results Table, Chunk Summary Table, and rose diagram for this data set. This functionality allows users to quickly and easily calculate a comprehensive PCP data set for all sample micrographs of an experimental group taken from the same region. Second, the PCPA plug-in suite also contains a stand-alone rose diagram feature which allows users to input any numerical angle measurement dataset to obtain descriptive circular statistics and a rose diagram, regardless of whether or not the data set was collected via PCPA. Step-by-step instructions for using batch processing and the standalone rose diagram function can be found in the PCPA user manual.

### PCPA reliably recognizes and measures directionality in cochlear hair cells

To test PCPA’s ability to count cells and calculate PCP we first tested PCPA on mouse cochlear hair cells. Two investigators were provided with 24 cochlear images from P4 mice (2 images per cochlear apex, middle, and base; n = 4 mice). Each investigator independently preprocessed the images and ran PCPA. Angle calculations from PCPA were rated for accuracy by a third investigator using a 1-5 agreement score (Fig 2) and overall accuracy scores per cochlear turn were calculated for each rater (Table 1). For cell counting, PCPA achieved 99.81% accuracy with PCPA being able to readily detect 1,538 out of 1,541 hair cells across the images analyzed. For PCP angle measurements, 96.35% (apex), 97.81% (middle), and 98.06% (base) of cells analyzed were scored as a 1 (perfect measurement). When cells scored 1 and 2 were combined—indicating cells that had <10° variation between PCPA measurements and idealized retrospective manual measurements—99.41% (apex), 98.93% (middle), and 100% (base) of cells analyzed by PCPA met this criterion. These data suggest that PCPA is able to accurately measure over 96% of βII-spectrin labeled hair cells in a data set, with perfect precision as determined by a human rater, and this metric jumps to over 98% when allowing for a <10° deviation from ideal.

**Fig 2.**
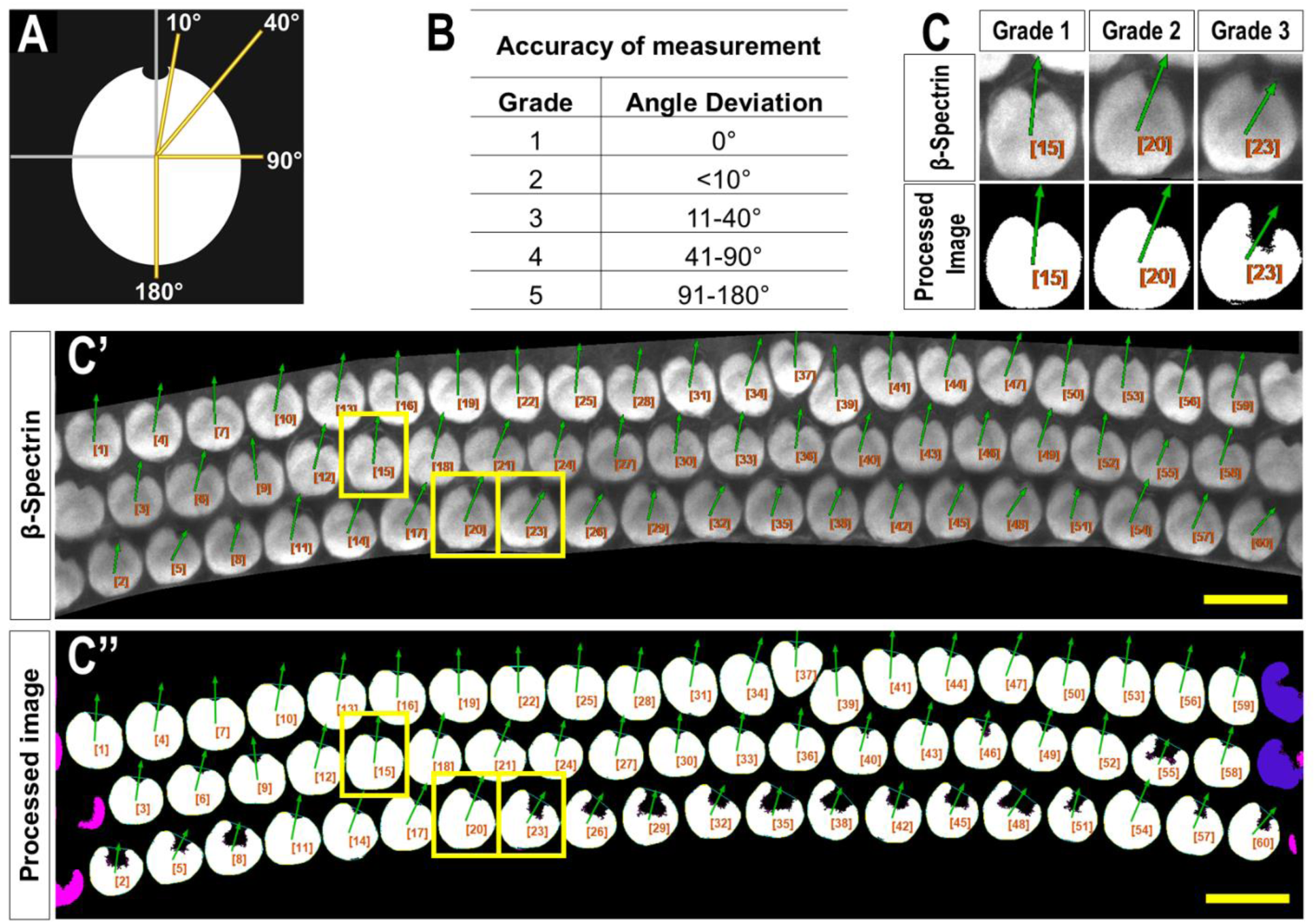
Criteria used to evaluate PCPA angle measurement accuracy. **(A)** A blinded investigator was instructed to consider an accurate planar cell polarity measurement as a ray originating from the center of mass of a cell and extending through the center of the fonticulus. The investigator then evaluated the directional arrow drawn by PCPA and assigned each cell an accuracy score from 1-5 using the grades shown in the table in **(B)** and the schematic from (A) as a reference. **(C)** Magnified view of example cells matching angle accuracy scores of 1, 2, and 3 from a **(C’’)** representative image of cochlear hair cells; yellow boxes indicate cells chosen for magnification, size excluded cells are pseudo-colored pink, and border excluded cells are pseudo-colored blue. Scores beyond 3 were not reported for any of the images analyzed (scale bars = 10 µm).

**Table 1.**
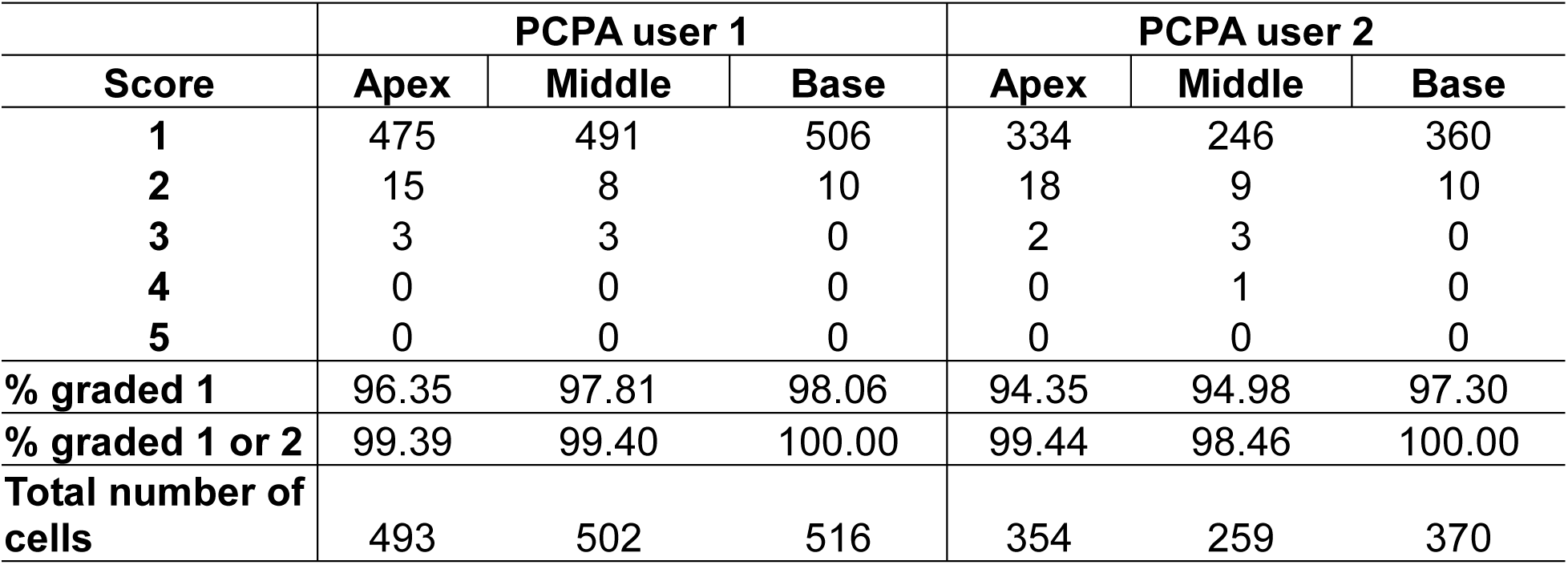
Accuracy of PCPA angle measurements from P4 cochleae.

Though βII-spectrin staining of the apical surfaces of hair cells presents as rather ideal circular chunks and caves, we tested whether PCPA could be used with another commonly used fluorescent label: phalloidin. Phalloidin binds to the F-actin rich hair cell stereocilia and the vertex of the “V” shape of the stereocilia bundles in cochlear outer hair cells has been used to assess cell polarity [29–31]. After preprocessing and making the phalloidin images binary, we found that application of the Plastic Wrap function allows PCPA to bridge the distance between the legs of the phalloidin positive bundle, and thus create a large cave. Selecting the largest cave as the directional point of interest causes PCPA to calculate polarity angles that were directionally opposite to measurements using the fonticulus as the directional point of interest (Fig 3A-3B). We compared PCP measurements obtained from βII-spectrin to the reversed values of those obtained from phalloidin on the same samples (acquired at the same time as multi-channel images that had high SNR for both phalloidin and βII-spectrin). When the phalloidin imaging was optimal, PCPA was able to measure the same number of outer hair cells with similar angle results (Fig 3A-C). Across 8 images of outer hair cells from N = 3 mice PCPA results indicated that the phalloidin approach resulted in fewer hair cells measured (n = 385) compared to the βII-spectrin approach (n = 428). Mean angle measurements remained similar, with a mean angle measurement of 79.997° (18.215°) for the phalloidin staining approach and a mean angle measurement of 80.174° (9.605°) for the βII-spectrin staining approach, though it should be noted that the variability was greater for the phalloidin images (Fig 3E and 3F). These results show the capability of PCPA to perform automated analysis of cell polarizations and to obtain reliable mean angle measurements from multiple markers. Although the data suggest that using βII-spectrin staining results in slightly greater numbers of cells being measured and decreased variability compared to phalloidin, we propose that these discrepancies reflect the reduced reliability of phalloidin as a marker compared to spectrin, pericentrin, and other antibody labeling that clearly demarcate the position of the kinocilium or its basal body, which has been suggested previously [32,33].

**Fig 3.**
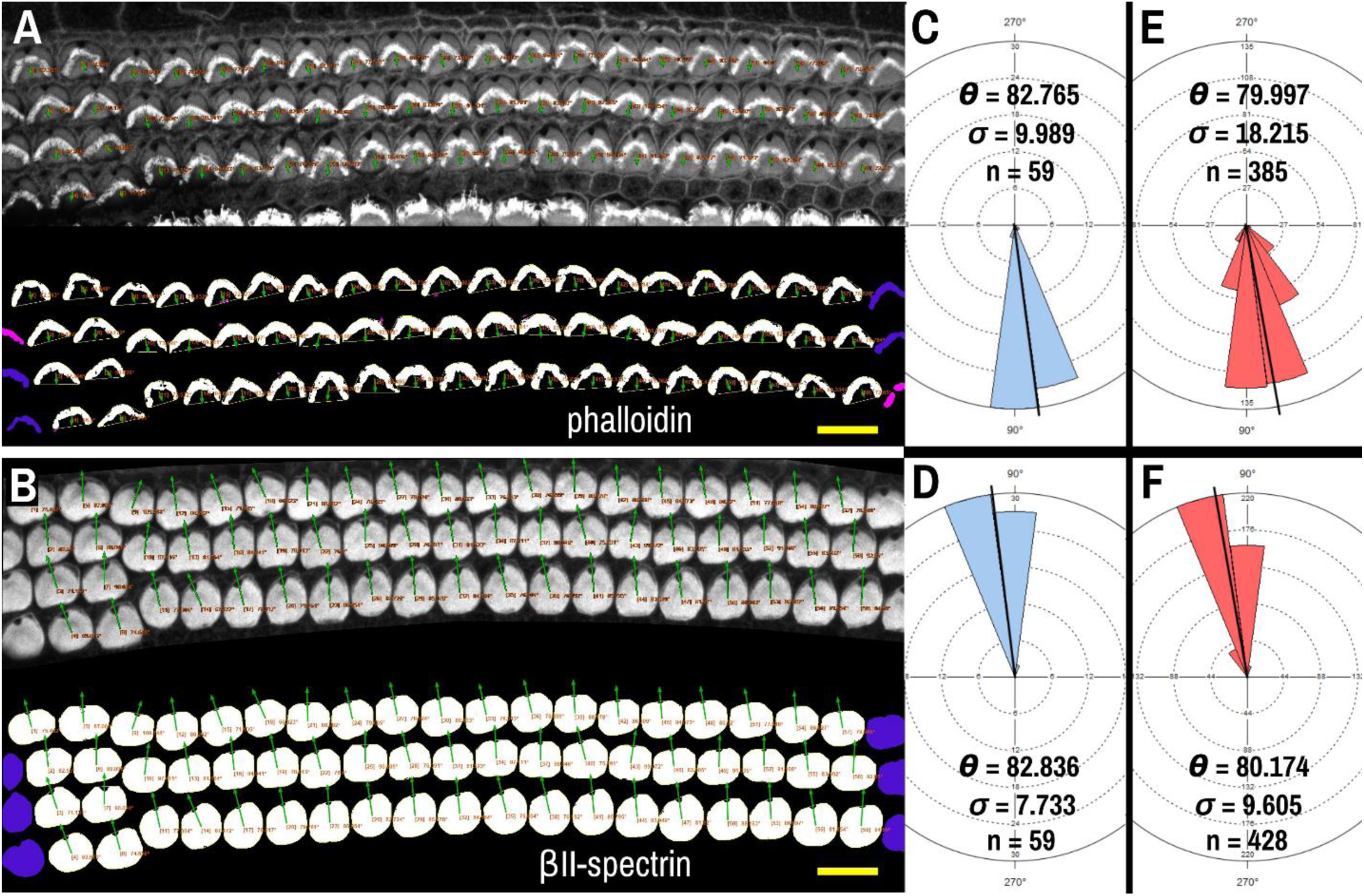
PCPA analysis of cochlear outer hair cells labeled with AlexaFluor488 conjugated phalloidin. Phalloidin **(A)** and βII-spectrin **(B)** labeled outer hair cells were imaged in different channels of the same P4 mouse cochleae, then the images were separated and thresholded for PCPA analysis (thresholded outputs shown in the bottom half of each panel). In both (A) and (B), size excluded cells are pseudo-colored pink and border excluded cells are pseudo-colored blue. **(C)** PCPA analysis of the image in (A) resulted in angle measurements collected from n = 59 outer hair cells with a mean angle (θ) of 82.765° and a standard deviation (σ) of 9.989°. **(D)** PCPA analysis of the βII-spectrin channel from this same cochlear region also measured angles from n = 59 outer hair cells with a resultant mean angle of 82.836° and a standard deviation of 7.733°. **(E)** The analysis was expanded to 8 images of phalloidin-labeled cochlear outer hair cells; PCPA was able to measure n = 385 cells and arrive at a mean angle of 79.997° (σ = 18.215). **(F)** PCPA analysis of βII-spectrin labeled cells from the same 8 images resulted in angles being measured from n = 428 outer hair cells with a mean angle of 80.174° (σ = 9.605). Thus, PCPA is able to measure planar cell polarity using images of phalloidin labeled cochlear outer hair cells. However, while the resultant mean angles in (E) and (F) were highly similar, the number of cells measured was greater when βII-spectrin labeling was used, and the variability of the measurements was reduced as compared to the phalloidin data (F vs. E). (scale bars = 10 µm).

### PCPA angle measurements are comparable to manually derived angle measurements in the utricle

The performance of PCPA was next tested on E17.5 utricle images which had been collected as control samples for a separate project in the laboratory. As these images were collected prior to the development of PCPA, the image acquisition parameters were not optimized to maximize βII-Spectrin fluorescent intensity, thus making them ideal “real-world” examples for testing. Sample boxes (500 x 500 pixels) were taken from posterior lateral extrastriolar, central lateral extrastriolar, anterior lateral extrastriolar, striola, posterior medial extrastriolar, and anterior medial extrastriolar regions (Fig 4). Cell angle measurements were collected manually by investigators instructed to bisect the hair cell apical surface and the fonticulus using the Fiji arrow tool and to record the angle as measured by Fiji. Another investigator preprocessed the images, converted them to binary, and used PCPA to measure the angles of the same cells that were quantified manually. Since the manual measurements did not indicate which angle measurement corresponded to which individual cell in any given image, the degree of agreement between the two data sets was assessed using the mean angle value from each image or image-pair as shown in Figure 5A-5L (n = 54 mean angle measurements from 5 separate embryos from 2 distinct litters). Average angle quantified per region was comparable between manual quantification and PCPA. The data are reported in Table 2. Calculation of the concordance correlation coefficient (CCC) showed a high level of agreement between the manually quantified data and PCPA quantified data (CCC = 0.999; 95% C.I. [0.9983, 0.9994]) (Fig 5M). Calculation of the Bland-Altman limits of agreement suggest minimal bias for the PCPA measurements as compared to manual measures, with a value of 0.482 degrees (95% C.I. [-0.676, 1.64]). The limits of agreement (LoA) were calculated at a 95% agreement level with the lower LoA at −7.833 (90% C.I [-9.143, −6.523]) and the upper LoA at 8.798 (90% C.I. [7.487, 10.108]). As shown in Figure 5N, the Bland-Altman analysis suggests that nearly all PCPA measured mean angles from similar studies could be expected to fall between −9 and +10 degrees of manually obtained mean angle measures.

**Fig 4.**
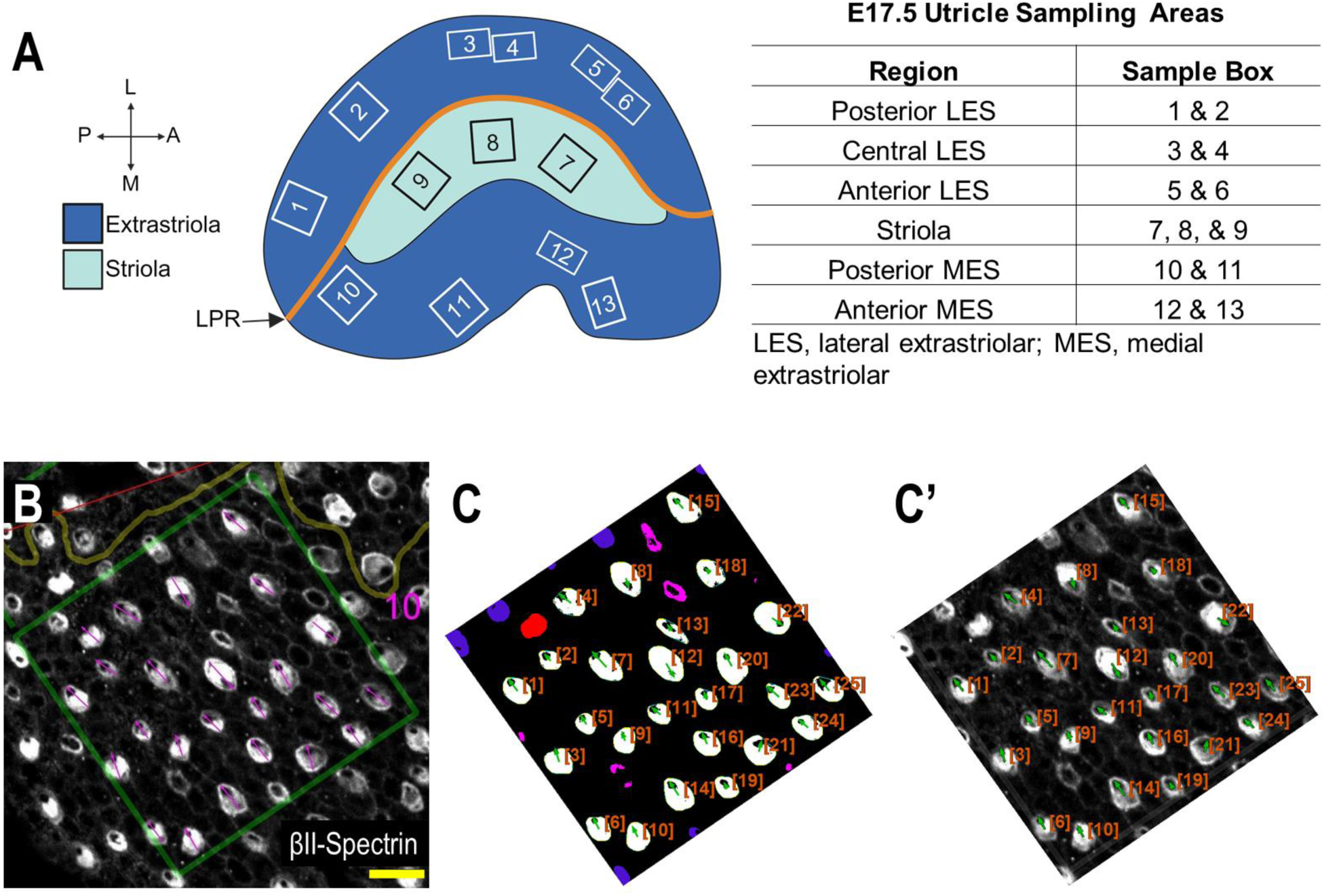
Performance of PCPA against manual PCP measurements for the E17.5 utricle (Fig 4A created with BioRender.com). **(A)** Example of regional sampling for E17.5 utricle analysis. Boxes were generally aligned according to the line of polarity reversal (LPR). **(B)** Example of manually collected PCP measurements from the box 10 region of a E17.5 utricle. **(C)** The image from (B) was preprocessed, made binary, and ran through PCPA analysis. PCPA angle measurements show high agreement with manual measurements in (B). **(C’)** Directional arrows (green) and cell ID overlays (orange) were combined with the original micrograph. (scale bars = 10 µm).

**Fig 5.**
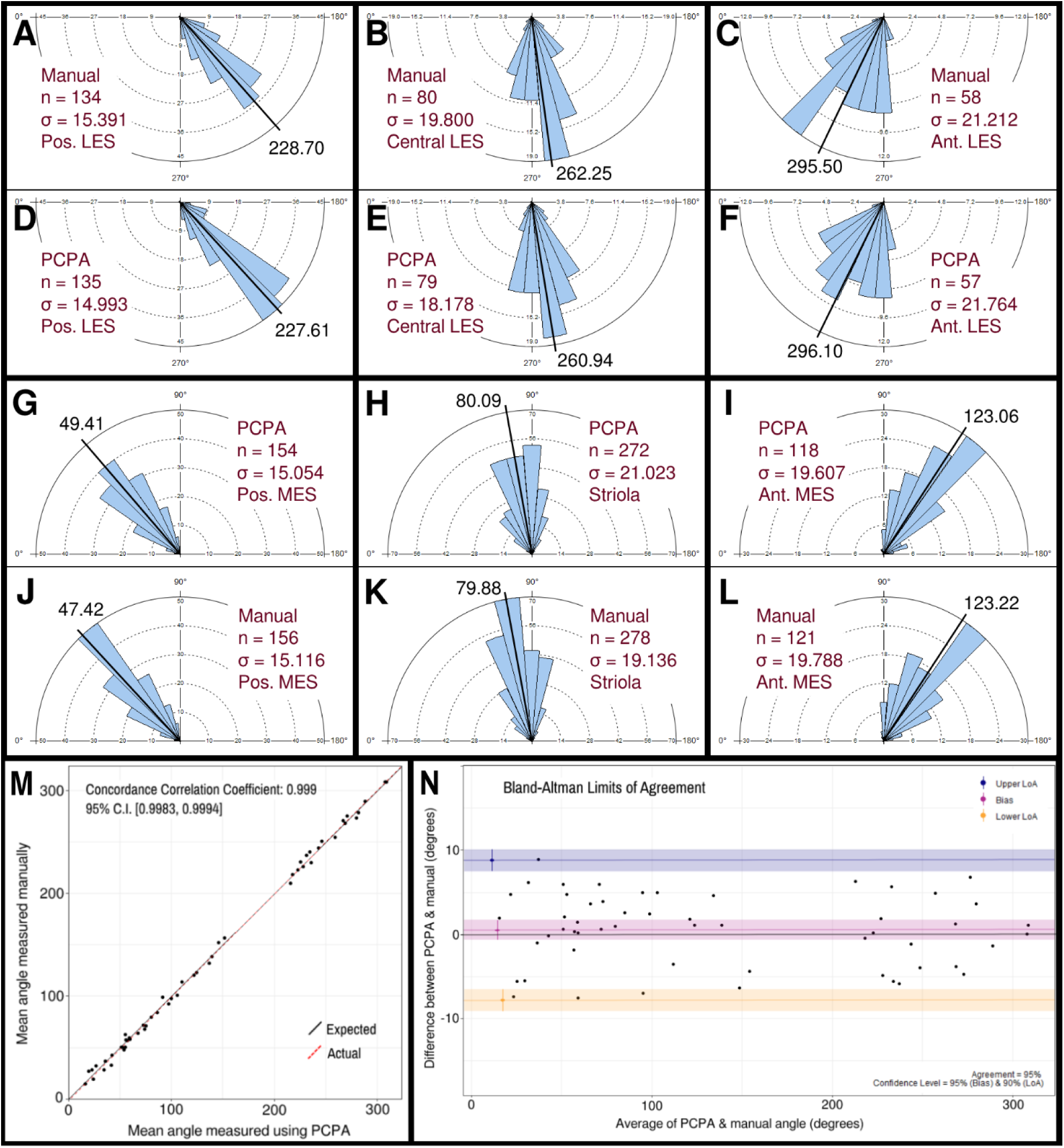
Manual vs. PCPA obtained angle measurements of βII-spectrin labeled hair cells in E17.5 mouse utricles. Manual measurements of hair cell orientations with respect to the LPR were taken as described in the manuscript main text and in Fig 4. A Rose diagram was plotted using PCPA (or stand-alone Rose Diagram plug-in) to visualize the results from each region investigated. For each region, the mean angle is represented by a black line and the exact value reported in black text. The number of cells in each region that were measured (n) and the circular variance in degrees (σ) is also provided. Manually obtained data for the LES are presented in the outer rows and can be directly compared to PCPA data in the central rows such that manual vs PCPA for the posterior LES region (A vs D), central LES (B vs E), and anterior LES (C vs F) are readily visualizable in the top two rows and manual vs PCPA data for the posterior MES (J vs G), striola (K vs H) and anterior MES (L vs I) are visualizable in the fourth and third rows. (M) Inter-rater reliability was assessed for manual vs PCPA measurements by plotting the mean angles for each image as a function of the PCPA value along the x-axis and the manual value along the y-axis, which, in the case of perfect agreement would yield a bisecting diagonal line (“Expected” shown in black). A line of best fit for the actual values (“Actual” shown in red) was plotted and the concordance correlation coefficient calculated as 0.999 (where a value of 1 would indicate absolute agreement for all values). (N) A Bland-Altman plot was generated again using the mean angle values from each image analyzed and upper and lower limits of agreement were calculated. The red line suggests a very small value for bias (<0.5 degrees) with a confidence interval (pink) that lies on both the positive and negative sides of zero. The blue and yellow lines represent the upper and lower limits of agreement, respectively, at a confidence level of 95%.

**Table 2.** Utricle measurements: Manual quantification vs. PCPA.

|  | Manual quantification |  |  | PCPA |  |  |
| --- | --- | --- | --- | --- | --- | --- |
| Region | N | Mean angle | SD | N | Mean angle | SD |
| Posterior LES | 134 | 228.70° | 15.391° | 135 | 227.61° | 14.993° |
| Central LES | 80 | 262.25° | 19.800° | 79 | 260.94° | 18.178° |
| Anterior LES | 58 | 295.50° | 21.212° | 57 | 296.10° | 21.764° |
| Posterior MES | 154 | 49.41° | 15.054° | 156 | 47.42° | 15.116° |
| Striola | 272 | 80.09° | 21.023° | 278 | 79.88° | 19.136° |
| Anterior MES | 118 | 123.06° | 19.607° | 121 | 123.22° | 19.788° |

LES, lateral extrastriolar; MES, medial extrastriolar

### PCPA automated measurement of polarity in a variety of tissues and from different species

Because planar cell polarity is found in a variety of tissues outside of the inner ear, we sought to test the PCPA plug-in on other image types including those using different labels, as well as samples from both wild-type animals and PCP mutants. To accomplish this, we collected images from several peer reviewed publications, preprocessed the images, ran PCPA, then compared PCPA’s results to published results. We first tested PCPA using images of radial glial cells from E16 and E18 mice from Mirzadeh et al. [34] In that report, cell junctions were labeled with an anti-beta-catenin antibody and primary cilia were labeled using an anti-γ-tubulin antibody (red and green, respectively, in the published figures). PCPA results (Fig 6) demonstrate the effective measurement of cell polarity with a decrease in variability in polarization evident from E16 (SD = 71.1) to E18 (SD = 56.5), consistent with the increasing alignment of the cells with developmental progression.

**Fig 6.**
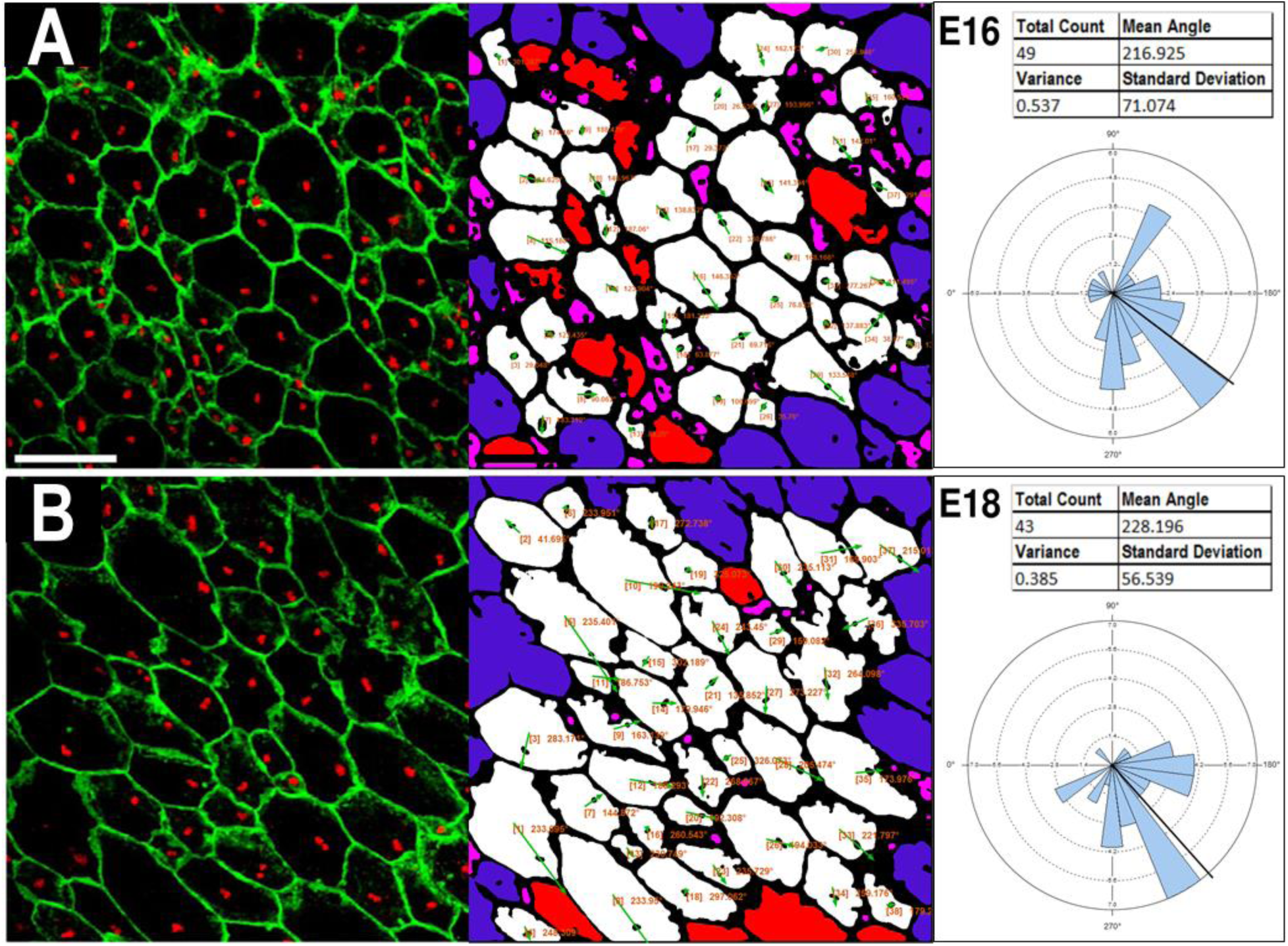
PCPA effectively measures radial glial progenitor cell orientations. Images reproduced from Mirzadeh et al. [34] demonstrate anti-β-catenin cell junction labeling (green) and anti-*γ*-tubulin immunolabeling of primary cilia (red) in radial glial progenitors from **(A)** E16 and **(B)** E18 mouse embryos. PCPA analysis of these images correctly defines the predominant angle as toward the bottom left corner of the images (216° to 228°) and demonstrates reduced variability as the alignment of cell direction becomes more consolidated during normal development from E16-E18.

Next, we tested PCPA’s performance on ependymal cell orientation using images from wildtype and *Celsr1* knockout mice (S5; Fig 7). In the original report, Boutin et al. [35] manually traced the perimeter of each ependymal cell using anti-ZO1 labeled cell junctions, then traced the perimeter of the cell’s γ-tubulin+ cilia patch. They then determined cell directionality utilizing a custom MATLAB script that, similar to PCPA’s approach, calculated angles based on the center of mass of the traced perimeter of the cilia patch in relation to the center of mass of the traced perimeter of the cell junction. We compared cell orientation results reported in Boutin et al. [35] to those generated by PCPA and found that PCPA was able to replicate Boutin et al.’s [35] angle orientations with a high degree of agreement (Fig 7; S5). One noted exception occurred in the *Celsr1* knockout sample where one cell’s PCPA-derived angle projection was nearly 180° opposite to the published data (cell number 6; Fig 7C-7C’). Because Boutin et al.’s [35] preprocessing workflow required an investigator to manually trace the cell perimeter, we speculated that the difference between PCPA and the published result could be due to the tracing process. Indeed, when we re-ran PCPA using Boutin et al.’s [35] tracings (Fig 7D-7D’), the directional disagreement for cell number 6 was resolved.

**Fig 7.**
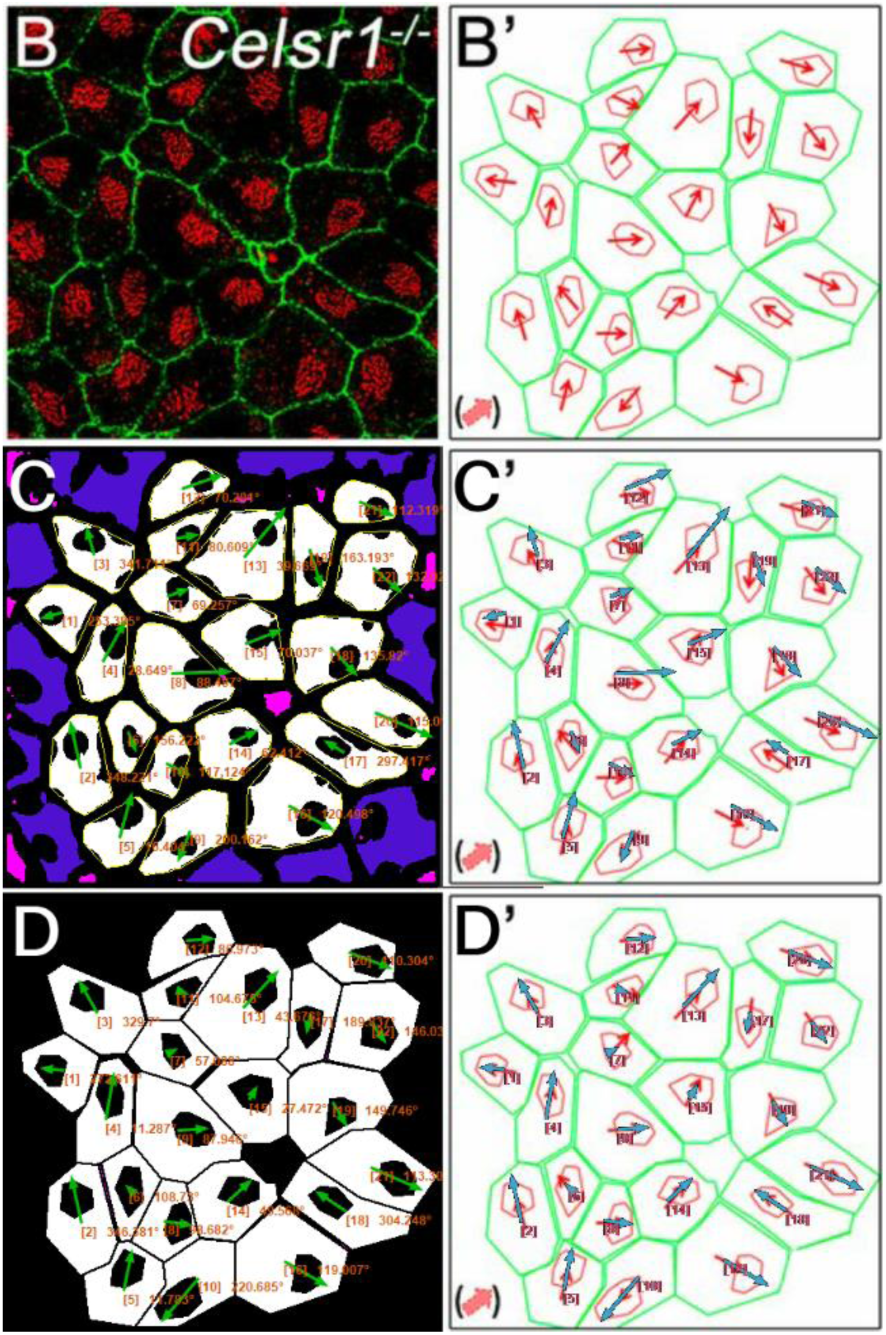
PCPA recapitulates angle measurements from ependymal cells in a Celsr1 knockout mouse model. Panels (B) and (B’) are taken from Figure 4 of Boutin et al. [35]. **(B)** A confocal image of Celsr1^-/-^ mouse ependymal cells labeled with antibodies against ZO-1 (green) and γ-tubulin (red). **(B’)** Manual tracings (green and red) of the cells in (B) with overlaid angle measurement arrows (red), as originally shown in Boutin et al. [35]. **(C)** Annotated output of image (B) after preprocessing and data collection with PCPA. The annotated output shows Plastic Wrap (yellow), edge excluded chunks (blue), size excluded chunks (pink), cell ID number with angle measurement (orange text), and vector arrows (green). **(C’)** Arrows from PCPA analysis (blue) are overlaid onto the image from (B’) to show the high level of agreement between PCPA and Boutin et al.’s [35] published data. Only one cell exhibited stark disagreement (cell #6), where the angle measured by PCPA appears nearly 180° opposite the published result. **(D)** Since Boutin et al. [35] measured manual tracings of the original image, we preprocessed then ran the manual tracing image through PCPA. **(D’)** Overlaid arrows from PCPA analysis (blue) ran using manual tracings show complete agreement with those in the original published work, suggesting any deviations in C and C’ are likely the result of differences in pre-preprocessing approaches (i.e. the use of a thresholded image versus manual tracings).

Finally, we compared PCPA’s performance on brightfield images of osmium (OsO_4_) fixed *Drosophila* ommatidia from Koca et al. [36]. Analysis of Drosophila ommatidia required a slight alteration to the preprocessing steps established above for PCPA to isolate the rhabdomeres in the photoreceptor cells (see methods section for details). PCPA produced angle orientations comparable to the manually obtained orientations collected by Koca et al. [36] for ommatidia from WT (Fig 8F-8H) and *Abl*-overexpression mutants (Fig 8I-8K). PCPA measurements from these images have near complete agreement to the published results (Fig 8H and 8K), and demonstrate the ability of PCPA to detect disrupted cell polarization in PCP mutants (Fig 8D vs 8E and 8H vs 8K). All together, these results show that PCPA has the ability to collect PCP angle measurements from a variety of diverse cell and tissue types collected with various staining and imaging parameters, and works in models with normal or disrupted PCP.

**Fig 8.**
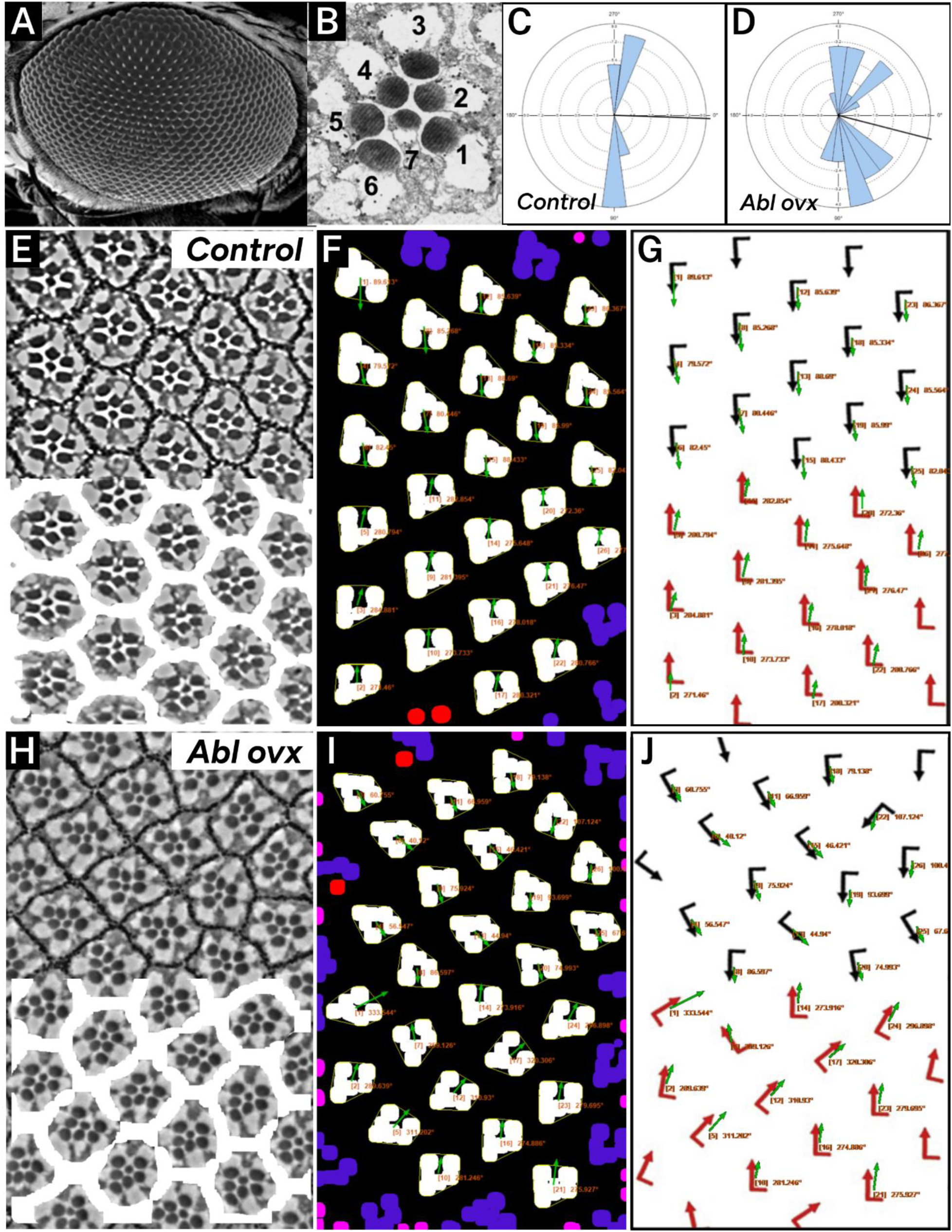
PCPA replicates angle measurements of wildtype and Abl overexpressing *Drosophila* ommatidia. Panels (A-B), derived from Tomlinson et al. [51] show the eye of the fruit fly at **(A)** low power under a scanning electron microscope and **(B)** after sectioning and at higher power under a light microscope to highlight the 7 readily visible rhabdomeres from which PCP can be determined. **(C)** A rose diagram generated by PCPA of wildtype/control Drosophila ommatidia (shown in panels E-G) shows that the ommatidia are well aligned along the vertical axis. **(D)** A rose diagram generated by PCPA of Abl overexpression in Drosophila ommatidia (shown in panels H-J) shows disruption of the vertical alignment. Images for analysis were copied from Koca et al. [36]. **(E)** Wildtype/control ommatidia (top) with overlay mask (bottom) created during PCPA preprocessing to mask ommatidial junctions. **(F)** PCPA annotated output showing Plastic Wrap (yellow), edge excluded chunks (blue), size excluded chunks (pink), unmeasurable chunks (red), the resulting angle measurements (orange text), and vector arrows (green). **(G)** The PCPA output overlay (green arrows) was added to the measurements presented in Koca et al. [36] (red arrows) to show the general agreement between PCPA and published measurements. **(H)** Abl overexpressing ommatidia (top) with overlay mask (bottom) created during PCPA preprocessing to mask ommatidial junctions. **(I)** PCPA annotated output showing Plastic Wrap (yellow), edge excluded chunks (blue), size excluded chunks (pink), unmeasurable chunks (red), the cell ID number with resulting angle measurements (orange text), and vector arrows (green). **(J)** The PCPA output overlay (green arrows) was added to the measurements presented in Koca et al. [36] (red arrows) to show the general agreement between PCPA and published measurements for the more disorganized ommatidia of the Abl overexpression mutant.

## Discussion

Numerous fields in biology rely on the quantification of cell numbers to assess proliferation, survival, and differentiation of cell populations. The process of counting cells manually is tedious and time consuming. Quantification of other cell characteristics, such as PCP, are equally important for answering questions related to developmental biology, and require greater efforts to manually quantify. As such, an automated method to assess micrographs and return accurate cell counts and angle measurements represents an important advance that should greatly benefit many disciplines in biology. In this study we present PCPA, a novel Fiji-based plug-in suite that automates the process of cell counting and calculation of PCP angle measurements. This user-friendly plug-in provides easily visualizable outputs (including rose diagrams, image overlays, and summary statistics) and has the ability to greatly speed up data collection for cell counting and PCP measurements while reducing potential bias or error in datasets.

We tested PCPA’s ability to collect cell counts and PCP angle measurements from βII-spectrin labeled cochlear and vestibular hair cells. Retrospective grading of the cochlear PCPA analyses and a direct head-to-head comparison between PCPA and manual measurements from vestibular hair cells showed that PCPA was able to accurately count cells present in the images, and measure the angles of orientation of those cells. While the main advantage of PCPA lies in the massive time-savings it can provide for researchers, it is also reasonable to hope that the automation of data collection will overcome any data biases introduced by human measurements. We acknowledge, however, that the preprocessing steps recommended to make high quality binary images for use with PCPA still rely upon the manual adjustment of micrograph images and could be a place where bias could be introduced. To combat this, we first suggest that investigators acquire images with homogeneous and high SNR, thus minimizing the need for pre-processing and allowing for potential automation of thresholding images to binary in Fiji (“Process > Binary > Make binary…” or other options such as Auto Threshold). However, consistently obtaining high quality micrographs, with high SNR, especially across different samples, is often challenging. Indeed, this is a limitation inherent in several previously published cell quantification programs as well [26,37,38]. While we were able to successfully run PCPA on images of suboptimal quality (both in terms of image dpi and antibody SNR ratio) by using the pre-processing steps recommended, we have also included several features in the PCPA outputs (i.e. overlays of arrows, cell numbers, and angle measurements) which are easily savable. These outputs can then be checked by experimenters, impartial observers, or any others persons wishing to verify the validity of the data.

In this report, we directly tested PCPA on samples from mouse inner ears, but also demonstrated the versatility and applicability of PCPA on a variety of cell morphologies and structures by comparing PCPA’s output to previously published studies examining PCP in murine radial glia [34], murine ependymal cells [35], and *Drosophila* ommatidia [36]. While minor changes to image preprocessing approaches were sometimes required in order to conform to PCPA’s central requirement of chunks being represented as white pixels and caves being represented as black pixels, PCPA showed robust ability to replicate PCP measurements from these diverse cell types.

Since the role of gene mutations in disruption of PCP phenotypes is a popular avenue of research in developmental biology, it was necessary to test the ability of PCPA to collect accurate data from sample images containing a wide degree of cell orientations. To do so, we tested PCPA on images along the line of polarity reversal (LPR) from E17.5 mouse utricles. The LPR is the thin zone spanning the antero-posterior axis of the utricle along which all hair cells orient; cells located on opposite sides of the LPR will exhibit extreme changes in orientation, on the order of approximately 180° reversals. Similar images with directly opposing orientations came from *Drosophila* ommatidia and again, PCPA’s angle measurements agreed with manual angle measurements, which highlights the capability of PCPA to obtain accurate measures from images with disorganized and/or highly varied cellular patterning. We further validated PCPA’s ability to measure cells with variable orientations by running PCPA on images from previously published papers featuring PCP disruption. PCPA performed similarly to the results published in Boutin et al. [35] to detect changes in ependymal cell orientation in Celsr1 knockout mice. For Abl-overexpressing *Drosophila* ommatidia, PCPA was again able to replicate angle orientations nearly identical to those published in Koca et al [36].

Other research groups have published automated or semi-automated methods for counting cells and for collecting PCP measurements. Several of these approaches have been published or made available as uncompiled scripts, thus requiring users to have the means and ability to compile these scripts on their systems, and/or to have some degree of coding knowledge for debugging. Scripts developed by Siletti, Tarchini, and Hudspeth [39], and by Boutin et al [35], both require manual selection or tracing of the center and perimeter of each cell, which is time consuming and has the ability to introduce human error or bias. In comparison, our Jython based plug-in is installed and updated through the popular open source image processing software Fiji and features a guided user interface with highly customizable options for data collection to meet the specific needs of their cell population of interest. Furthermore, for most samples analyzed by PCPA, no manual tracing or drawing is applied by the user, thus making the analysis closer to the original raw data and eliminating one more step where bias or error could be introduced.

While other standalone programs or Fiji based applications aimed at automating PCP data collection have been developed, some have not been updated for many years or are linked to software programs that have been discontinued (e.g. Metamorph [34]). Some are only executable on certain operating systems (e.g. FijiWingsPolarity [40], which works with Mac OS, but not PC). Of those that remain, the underlying approaches to measuring cell orientation differ from PCPA. For example, QuantifyPolarity [41] and SEGGA [37] were developed to measure *Drosophila* cell wing polarity across time lapse images using fluorescence intensity gradients from developmental factors influencing PCP. These programs share some similarities with PCPA including preprocessing approaches that do not require manual drawing or segmentation, a graphical user interface, and user customization options. The main differentiating factor is the nature of the investigator’s approach to PCP visualization. QuantifyPolarity [41] and SEGGA [37] are more appropriate for approaches using fluorescently labeled proteins with a clear gradient across the cell or for time lapse imaging studies, while PCPA was designed for use with cell populations where the directional marker is a defined morphology, i.e. that of a chunk with a cave.

Another area in the field of automated image analysis that is gaining popularity at an exponential pace is artificial intelligence, or A. I. While we are not aware of reports of any currently available machine learning or A.I. workflows for the measurement of planar cell polarity, there are currently several published machine learning approaches for cell counting [42–44], and indeed several such approaches applied specifically to the counting of hair cells [38,45,46]. While these approaches are mostly successful in counting cochlear hair cells (or other cells of interest), machine learning approaches can face barrier-to-entry challenges to investigators not trained in computer sciences or in laboratories lacking the processing power required by some machine learning systems. A.I. approaches also often require training for each dependent measure (or combination thereof) for which they are to be used. In many cases, A.I. approaches require such training in each new laboratory due to differences in sample quality and imaging parameters. This extensive training has the potential to offset potential time savings, as users will have to make numerous manual measurements to provide the A.I. with feedback and validate its performance. A.I. approaches can also suffer from limited adaptability in the face of novel phenotypes or processing approaches not encountered during the training period [38]. Thus, while A.I. based approaches are likely to continue improving there is still a clear utility for a non-A.I. automated approach such as PCPA, which does not require training and retains a larger degree of user oversight and transparency.

Previously published automated cell counting programs also often struggle to cope with overlapping cells, which can require advanced machine learning systems [38], labor intensive optical clearing methods [45], or user guided segmentation of cells [26]. In the case of inner ear tissues this issue can be particularly problematic where, for example, there are little to no options for the labeling of cell nuclei in the neurons of the auditory nerve which makes segmentation nearly impossible. A similar problem persisted for decades with regard to sensory hair cells where myosin or other non-nuclear proteins were the best available markers. More recently, immunolabeling of the nuclear protein POU4F3 has been proposed in a semi-automated method for hair cell counting [26], but even with a nuclear marker, there appear to be difficulties with segmentation requiring manual separation of doublets and triplets of overlapping cells by the user. This problem may be partly solvable from the staining and imaging side of the approach where we have found that immunolabeling spectrin in hair cells provides better segmentation than staining of POU4F3 (data not shown).

However, even with βII-spectrin labeling, overlap events still occur. Our PCPA suite features some further ability to address these instances of merged cells, through the optional Doublet Splitting mode. By setting a maximum length: width ratio, users are able to target cell conglomerates that violate these parameters and split the aggregates in half, then subsequently treat each half of a split as an independent cell during further analyses. Doublet Splitting improves the ability to automate data collection in cell populations that experience occasional overlap of cell bodies, though this feature is limited to the ability to split one aggregate into halves and may not be useful for instances where cell density causes three or more cells to overlap. Still, users may alternatively choose to exclude cells by ratio or size, which is beneficial to avoid erroneous angle measurements, but is sub-optimal for cell counting purposes. However, PCPA color codes all excluded cells thus making it easy for a user to identify cell aggregates and manually adjust the final count. One further limitation of the doublet splitting function in PCPA is that doublet recognition is accomplished through setting a length vs width parameter relative to the image axis. Doublets that align more closely with vertical or horizontal axes are more likely to be split accurately down the middle of the doublet mass, and doublet splitting accuracy worsens as the alignment of a doublet approaches 45° relative to the image axis. Despite this limitation, Doublet Splitting was used quite effectively in our cochlear and vestibular samples, partially thanks to the low incidence of cell overlap afforded by the specificity of βII-Spectrin labeling of the apical portion of hair cells. Furthermore, the majority of doublets we encountered were aligned with the horizontal or vertical axis, allowing for a high rate of success of the Doublet Splitting feature.

Despite the listed caveats, the data suggest that the PCPA plug-in suite is a robust and accurate tool for the automated collection of cell counts and PCP angle measurements. Furthermore, the increased throughput provided by PCPA may lend it for use in phenotyping other processes beyond PCP. For example, perturbations in development, regeneration, cancer, or other processes, can result in tissue disorganization independent of the PCP pathway. The ability that PCPA provides to measure thousands of cells quickly could therefore suggest great enough power and sensitivity to detect more subtle disorganization of tissues as well as its potential use in higher throughput genetic or molecular screens. In summary, this plug-in suite was designed to be compatible with the widely used image processing software Fiji and features an easy to understand guided user interface. PCPA has been shown to perform comparably to traditional manual PCP measurement methods, and has the ability to greatly speed up PCP data collection while potentially reducing human error and bias in PCP datasets. PCPA has been shown here to be applicable to a number of cell and tissue types with varied cell morphologies, including murine inner ear hair cells, murine ependymal cells, murine radial glia, and *Drosophila* ommatidia. Finally, the PCPA plug-in has been developed with a good deal of assistive material for users. A detailed user manual for installing and using PCPA can be found at https://github.com/WaltersLabUMC/PCP-Auto-Count.git, and video tutorials demonstrating PCPA can be found online including a simple demonstration of pre-processing and analysis of a utricle sample at: https://www.youtube.com/watch?v=PTFlGv5Laa0.

## Materials and Methods

### Animal care

All mice used in this experiment were housed in a temperature-controlled animal vivarium under a 12:12 light/dark cycle with *ad libitum* access to standard chow and water. All procedures were approved by the University of Mississippi Medical Center Institutional Animal Care and Use Committee (IACUC) and followed the NIH guidelines for the care and use of laboratory animals.

To obtain utricles from embryonic day (E) 17.5 mice, a timed mating strategy was employed to cross *Six2(+/-)* female mice with *Six2(+/-)* males. The *Six2(+/-)* mice were created and maintained as a mixed background of C57Bl/6J and 129/Sv [47]. The male and female mice were paired in the evening, and on the following morning the male was removed and female mice were assumed pregnant with embryos at stage E0.5. On day E17.5, dams were euthanized and embryos were collected. The heads of the embryos were removed, rapidly hemi-sectioned, then placed into 4% paraformaldehyde for 4 hours at room temperature (RT) then stored in PBS at 4°C until use. Tail biopsies were collected into sterile microcentrifuge tubes for genotyping. Only wild type embryos were utilized for PCPA data collection. For postnatal cochlea experiments, CD1 mice were euthanized at postnatal day (P) 4 and temporal bones were collected into 4% paraformaldehyde for 4 hours at RT then stored in PBS at 4°C until use.

### Immunohistochemistry

E17.5 utricles were micro-dissected and collected for immunohistochemistry. Whole mount samples were washed in PBS then immersed in blocking buffer (0.2% Triton-X and 4% donkey serum) for 60 minutes at RT. Tissue was incubated in the following primary and secondary antibodies: βII-spectrin (1:200), Alexa Fluor 568 goat anti-mouse IgG1 (1:400), and Alexa Fluor™ 488 Phalloidin (1:800). Following a final PBS wash, samples were whole-mounted to slides and coverslipped with Fluoro Gel plus DABCO.

P4 cochlear whole mounts were washed in PBS then incubated in Image-iT™ FX signal enhancer for 30 minutes at RT. Following PBS washes, samples were immersed in blocking buffer (0.2% Triton-X and 4% donkey serum) for 60 minutes at RT. The Mouse on Mouse (M.O.M.) IgG Blocking Reagent kit was used per manufacturer’s instructions and the following primary and secondary antibodies were used: anti-Pou4f3 (1:200), anti-βII-spectrin (1:200), Alexa Fluor 647 goat anti-mouse IgG1 (1:1000), Alexa Fluor 568 goat anti-mouse IgG1 (1:1000), Alexa Fluor™ 488 Phalloidin (1:800), and Hoechst 33342 (1:1500). Following a final PBS wash, samples were mounted with Fluoro Gel with DABCO. A detailed list of all antibodies and reagents used, including catalog and RRID numbers is presented in Table S6.

### E17.5 utricle angle measurements

Z-stack images of the entire E17.5 utricular maculae were obtained by tile-scanning with a Zeiss LSM 880 confocal microscope (40X oil objective, 2048 x 2048 resolution). Maximum intensity projections were then created from the z-stack images and used in subsequent analysis. In order to systematically survey hair cells across different anatomical regions of the utricle, sample boxes (500 x 500 pixels) within the following regions were drawn (Fig 4): posterior lateral extrastriolar (boxes 1 and 2), central lateral extrastriolar (boxes 3 and 4), anterior lateral extrastriolar (boxes 5 and 6), striola (boxes 7, 8, and 9), posterior medial extrastriolar (boxes 10 and 11), and anterior medial extrastriolar (boxes 12 and 13). Boxes were oriented perpendicular to the line of polarity reversal as drawn onto the image by one of two investigators charged with manual angle measurements. To measure planar cell polarity, investigators were instructed to bisect the cuticular plate and continue through the center of the fonticulus, using the arrow tool in Fiji. Automated angle measurements were collected by a third investigator who was provided the same utricle images used for manual angle measurements described above. The images were converted to binary using the preprocessing steps outlined below, then PCPA was used to measure cell angles with the setting to detect the largest cave within each chunk.

### P4 cochlear hair cell angle measurements

Z-stack images of cochlear hair cells were taken from randomly selected areas from the apex, middle, and base of P4 cochleae using a Zeiss LSM 880 confocal microscope with a 63X oil immersion objective and 2048 x 2048 resolution. Two investigators independently preprocessed the cochlear images and ran PCPA to obtain angle measurements. Inner and outer hair cells were preprocessed and analyzed separately. One investigator used the setting indicating the largest fonticulus of a chunk as the cave of interest and the second investigator ran PCPA utilizing the setting indicating the northmost cave as the cave of interest. A third investigator was provided the original images with an overlay of arrows drawn by PCPA. This investigator assigned each analyzed cell an accuracy score from 1-5 (1 = perfect measurement; 2 = <10° deviation; 3 = 11°-40° deviation; 4 = 41°-90° deviation; 5 = 91°-180° deviation; Fig 2).

### Image preprocessing for PCPA

All preprocessing steps for cochlear and utricle images were conducted using in-built functionality of Fiji unless otherwise stated. Briefly, βII-spectrin or phalloidin staining were separated from multicolor images by splitting the channels (“Image > Color > Split channels”). To make the z-stacks two dimensional, maximum projections were made (“Image > Stacks > Z project > maximum”) and converted to 8-bit images (“Image > Type > 8 bit”). Next the “Subtract Background” function was applied (“Process > Subtract Background…;” rolling ball radius = 50 or 100 pixels depending on the user). Users manually adjusted the threshold value for each image (“Image > Adjust > Threshold”) with the goal of including as much cell fluorescence as possible while minimizing non-specific or background pixels. The “Process > Noise > Remove Outliers” function (radius = between 2-12 pixels; threshold = 50) was often employed to smooth the outer surfaces of chunks and reduce the risk of plastic wrap creating spurious small caves.

#### PCPA settings: P4 cochleae

All cochlear images were oriented such that the radial axis of the cochlea was vertical with inner hair cells toward the bottom of the image and outer hair cells toward the top. After each investigator conducted their own preprocessing, images were analyzed using the PCPA plug-in. Both investigators used the following PCPA settings: remove noise of ≤500 pixels or fewer, exclude chunks ≤10 pixels from the image border, apply the Plastic Wrap function, and separate doublets when chunks measured 1.5 times wider than they were tall. Investigators differed on the following settings: investigator 1 ignored caves ≤3 pixels and selected the largest cave as the directional point of interest. Investigator 2 ignored caves of ≤15 pixels and selected the northmost cave as the directional point of interest. Angle measurements were set on a counter-clockwise 0-360° scale where 90° pointed north.

#### PCPA settings: E17.5 utricles

After preprocessing, images were analyzed using the PCPA plug-in with the following settings: remove noise of ≤500 pixels or fewer, exclude chunks ≤10 pixels from the image border, apply the Plastic Wrap function, separate doublets when chunks measured 1.5 times wider than they were tall, ignore caves of ≤15 pixels and select the largest cave as the directional point of interest. Angle measurements were set on a counter-clockwise 0-360° scale where 90° pointed north.

#### Preprocessing and PCPA settings: previously published images

Images from published works were obtained by magnifying the online images in a browser then saving screenshots as tiff files. For two-color images, the channels were separated in Fiji and each channel underwent manual adjustment of threshold before re-merging of the two channels into a single binary image.

Images of murine radial glia from Mirzadeh et al. [34] and images of ependymal cells taken from Boutin et al. [35] underwent channel splitting (Image > Color > Split Channels), and each channel image was converted to 8-bit grayscale (“Image > Type > 8-bit”). Each channel image then underwent manual thresholding (“Image > Adjust > Threshold”). Noise removal (“Process > Noise > Remove Outliers;” radius = between 2-10 pixels; threshold = 50) was applied to each image and for the ependymal cells an additional binary erode step was taken (“Process > Binary > Erode”). Image channels were then re-merged (“Overlay > Add image…”), and the LUT was inverted (“Image > Color > Invert LUTs”).

Images of *Drosophila* ommatidia taken from Koca et al. [36] were each duplicated (Image > Duplicate), then a manual threshold was applied to the duplicated image to leave only the cell junctions visible, the LUT was inverted so that the junctional borders would appear white rather than black, and Fiji’s “Process > Binary > Dilate and/or Erode” was used to connect the lines as much as possible while leaving empty space in the center of each ommatidium. This image was then added back to the original as a zero-transparent overlay (“Image > Overlay > Add Image…;” select “Zero Transparent”) and the image was flattened (“Image > Overlay > Flatten”). This image was then manually thresholded (“Image > Adjust > Threshold”) to isolate the rhabdomeres. The LUT was then inverted (“Image > Color > Invert LUTs”), and binary dilate (“Process > Binary > Dilate”) was applied until rhabdomeres touched to form a u- or h-shaped chunk. PCPA was then run on the images using Plastic Wrap and the selection for the largest cave.

### Data analysis

Mean angles, resultant mean length (RML), circular variance, and circular standard deviation were calculated by PCPA using equations based on Fisher [48] (equations provided in the user manual). All rose diagrams and descriptive statistics generated using PCPA were checked for accuracy against rose diagrams and descriptive statistics generated from the same data using the circular package [49] in *R*. Inter-rater reliability for angular measurements of utricle hair cells was tested using the SimplyAgree [50] package in *R*. The mean of cell angle measurements from each image was paired between the manual blinded investigator and PCPA and from these a concordance correlation coefficient and Bland-Altman limits of agreement were generated, again using the SimplyAgree package in *R*.

## Author contributions

**Conceptualization:** Bradley J. Walters, Luke D. Baum.

**Data Curation:** Kendra L. Stansak, Bradley J. Walters.

**Formal Analysis:** Bradley J. Walters, Kendra L. Stansak.

**Funding Acquisition**: Bradley J. Walters

**Investigation:** Kendra L. Stansak, Sumana Ghosh, Punam Thapa, Vineel Vanga, Bradley J. Walters.

**Methodology:** Bradley J. Walters, Kendra L. Stansak, Sumana Ghosh.

**Project Administration:** Kendra L. Stansak, Bradley J. Walters

**Resources:** Bradley J. Walters, Sumana Ghosh, Punam Thapa, Kendra L. Stansak.

**Software:** Luke D. Baum, Kendra L. Stansak, Bradley J. Walters

**Supervision:** Bradley J. Walters.

**Validation:** Bradley J. Walters, Kendra L. Stansak, Vineel Vanga.

**Visualization:** Kendra L. Stansak, Bradley J. Walters.

**Writing – Original Draft Preparation:** Kendra L. Stansak, Bradley J. Walters.

**Writing – Review and Editing:** Kendra L. Stansak, Luke D. Baum, Punam Thapa, Bradley J. Walters.

## Acknowledgements

The authors would like to thank the staff of the UMMC Center for Comparative Research for their dedication to the care and welfare of experimental animals, Sylvia Kassoff for technical assistance, and the funding agencies noted above.

## Abbreviations

A.I.: artificial intelligence
CCC: concordance correlation coefficient
E: embryonic day
LPR: line of polarity reversal
P: post-natal day
PCP: planar cell polarity
PCPA: PCP-Auto Count plug-in
RML: resultant mean length
RT: room temperature
SNR: signal-to-noise ratio

