## Supporting figure S1 for "PCP Auto Count: A Novel Fiji/ImageJ plug-in for automated quantification of planar cell polarity and cell counting"

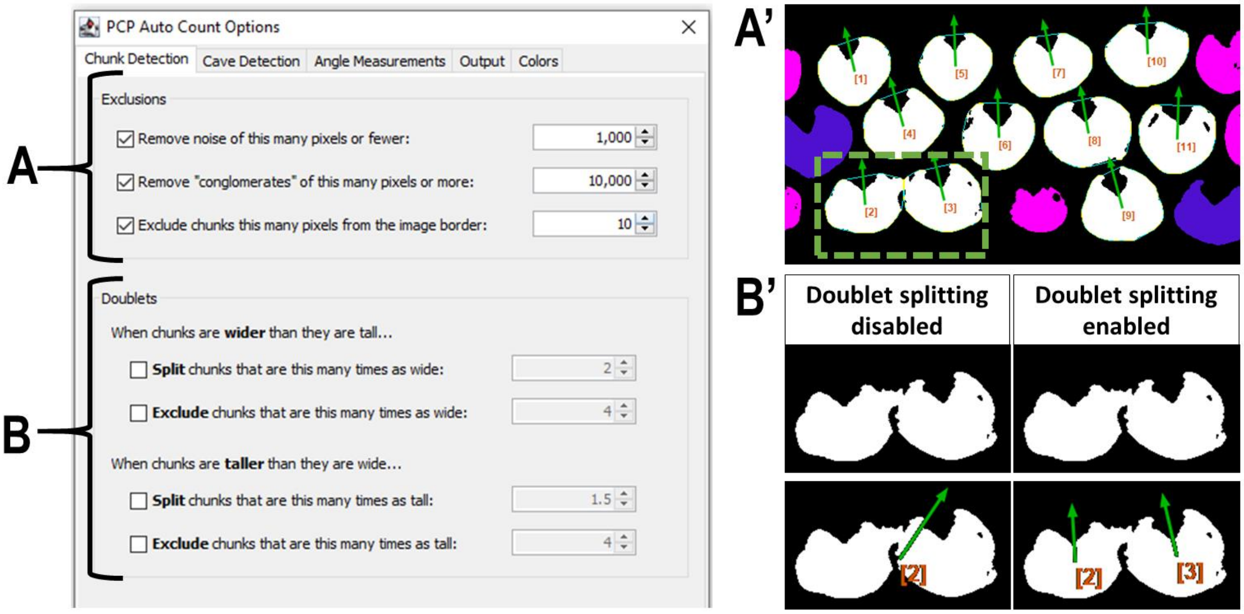


**S1.** User option menu for PCPA chunk detection. **(A)** The exclusion criteria box allows users to disregard chunks smaller and/or larger than a specified pixel size, and to exclude cells a set pixel distance from the image border. **(A')** Example output where size excluded chunks are shown as pink pseudo-coloring and border excluded chunks are shown as blue pseudo-coloring. **(B)** To address instances were two chunks touch after thresholding, users can set options to split chunks meeting a set ratio of wider-than-tall or taller-than-wide, or to exclude these chunks from further analyses. **(B')** Subset of A' (green box). If no doublet splitting options are applied, PCPA will treat this chunk as one mass and the angle measurements will incorrectly calculate based on the total chunk mass and dominant cave of the chunk. When doublet splitting options are applied, PCPA will split the doublet in half. The two resulting halves will be treated as independent chunks with chunk and cave centroid calculations being based on the mass contained within each respective box.
