## Supporting figure S2 for "PCP Auto Count: A Novel Fiji/ImageJ plug-in for automated quantification of planar cell polarity and cell counting"

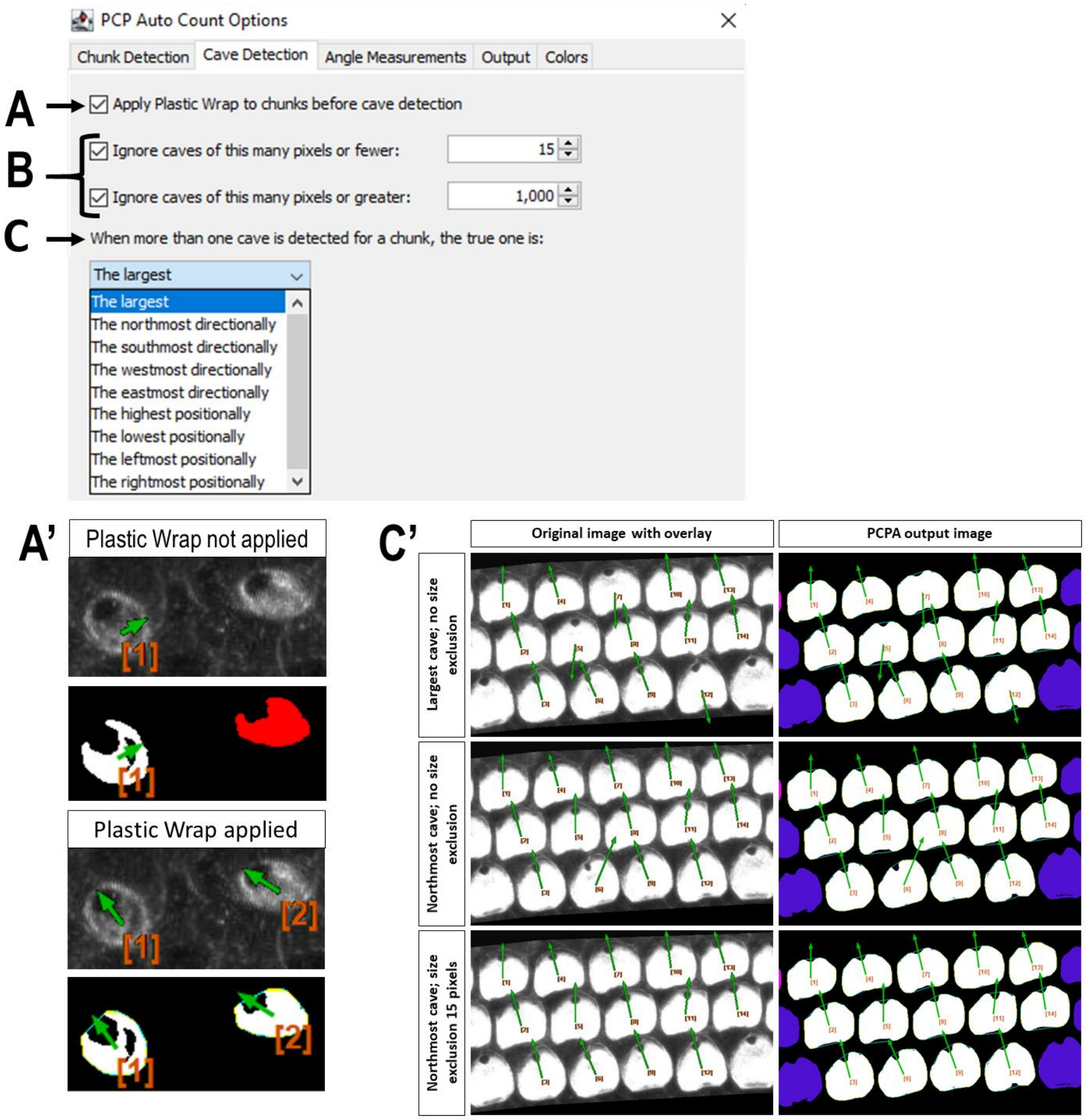


**S2.** User menu for PCPA cave detection. **(A)** Users are recommended to apply the Plastic Wrap function to their analyses. Plastic Wrap allows PCPA to group pixels that are proximal, but not abutting, into the same chunk. **(A')** Example images show that when plastic wrap is not applied, chunk 1 selects an incorrect cave, and chunk 2 is excluded from analysis for having no cave. When plastic wrap is applied, PCPA artificially closes off the open cave and selects the correct cave as the cave of interest. **(B)** Users can set minimum and maximum size requirements for cave of interest selection. **(C)** Users can set a directional requirement for cave of interest selection. Selecting a directional characteristic (e.g. northmost) will instruct PCPA to designate the cell's cave as the largest northmost inclusion. **(C')** Users can combine directional characteristics with a lower size limit for caves, in order to filter out incidental inclusions created by Plastic Wrap, pitting caused by the thresholding process, or large inclusions caused by irregular cell shapes.
