## Supporting figure S3 for "PCP Auto Count: A Novel Fiji/ImageJ plug-in for automated quantification of planar cell polarity and cell counting"

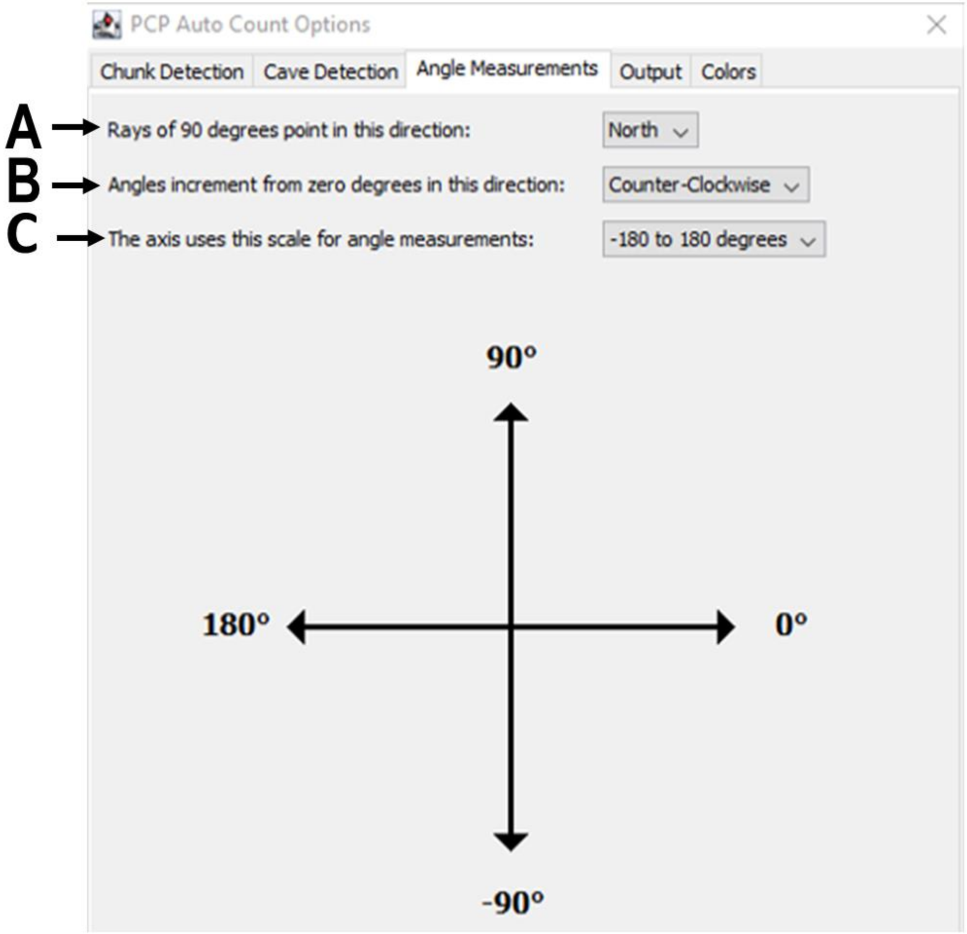


**S3.** User options for angle measurement calculations. **(A)** Option to set the direction of 90° ray. **(B)** Option to set direction of axis incrementation. **(C)** Option to set the scale of the (XY) axis to -180°/180° or 0°/360°
