## Supporting figure S4 for "PCP Auto Count: A Novel Fiji/ImageJ plug-in for automated quantification of planar cell polarity and cell counting"

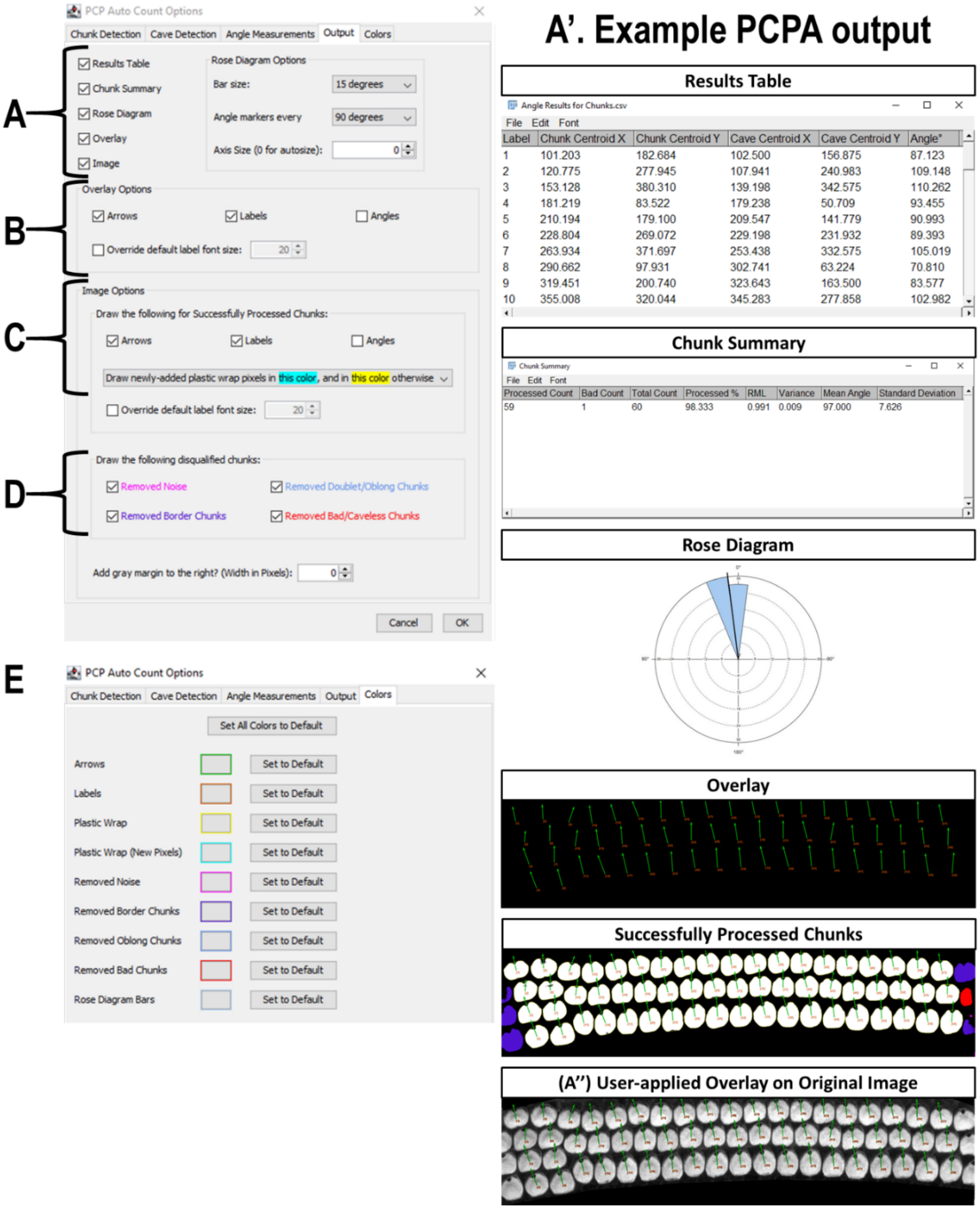


**S4.** PCPA data output tab. **(A)** Options for data metrics produced by PCPA. **(A')** Example of data metrics in A. **(A'')** Example of overlay superimposed on original preprocessed image. **(B)** The overlay (and selected overlay information) can be superimposed over the original image for data visualization. **(C)** Optional labels can be viewed on the post-PCPA image. **(D)** Color coding of the post-PCPA image can indicate to the investigator what chunks may have been excluded and why. **(E)** The Colors tab allows users to customize color coding options from D.
