## Supporting figure S5 for "PCP Auto Count: A Novel Fiji/ImageJ plug-in for automated quantification of planar cell polarity and cell counting"

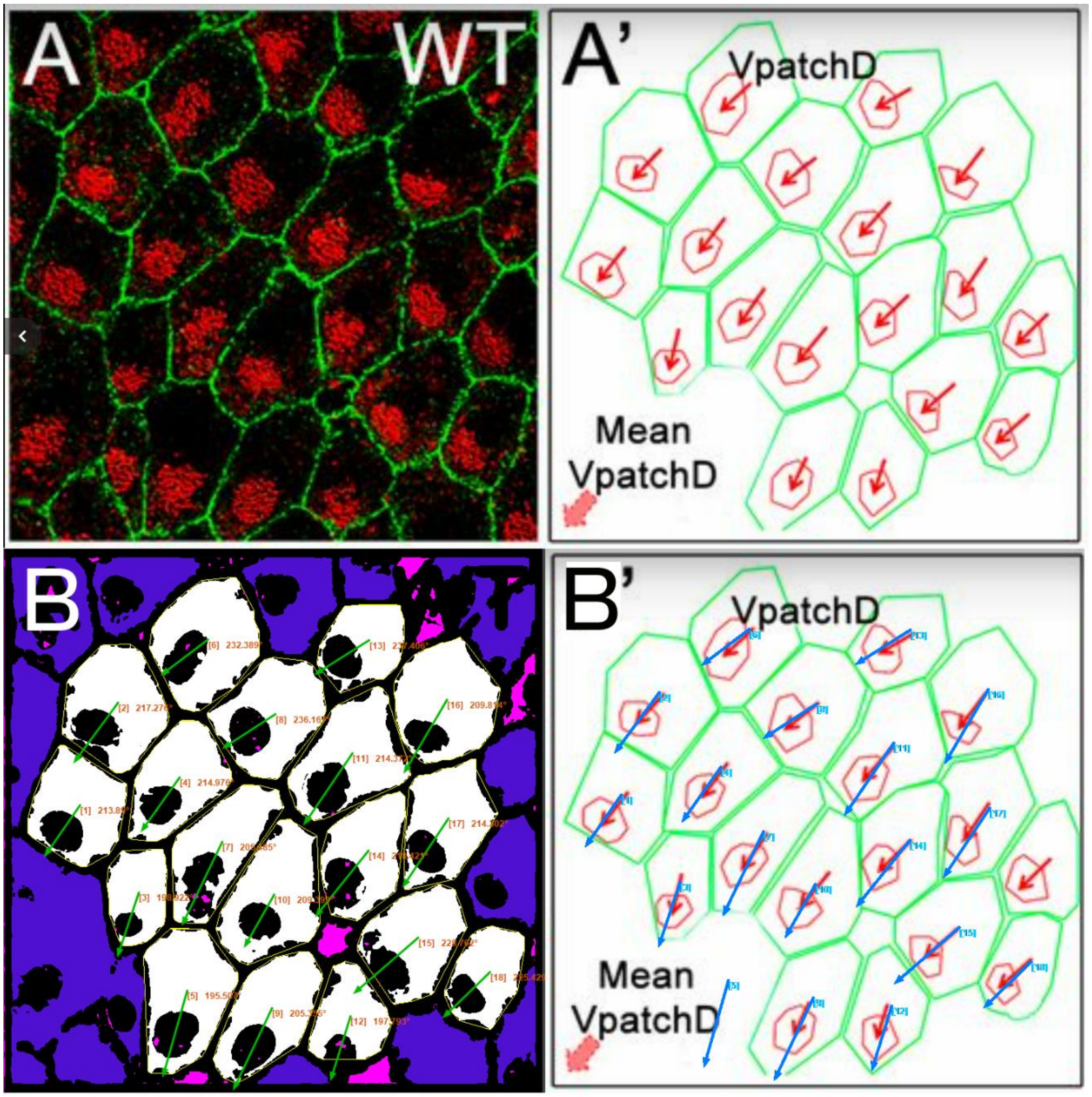


**S5. PCP Auto Count recapitulates angle measurements taken from WT murine ependymal cells**. Panels (A) and (A') are taken from Figure 4 of Boutin et al. [35]. **(A)** A confocal image of WT mouse ependymal cells labeled with antibodies against ZO-1 (green) and γ-tubulin (red). **(A')** Manual tracings (green and red) of the cells in (A) with overlaid angle measurement arrows (red), as originally shown in Boutin et al. [35]. **(B)** Annotated output of image (A) after preprocessing and data collection with PCPA. The annotated output shows Plastic Wrap (yellow), edge excluded chunks (blue), size excluded chunks (pink), cell ID number with angle measurement (orange text), and vector arrows (green). **(B')** Arrows from the PCPA analysis (blue) are overlaid onto the image from (A') to show the high level of agreement between PCPA and the angle measurements reported by Boutin et al. [35].
