## Supporting Table S6 for "PCP Auto Count: A Novel Fiji/ImageJ plug-in for automated quantification of planar cell polarity and cell counting"

**S6 Table**

| Materials and reagents used | | | |
|---|---|---|---|
| **Product** | **Company** | **Cat. number** | **RRID** |
| Anti-βII-spectrin | BD Biosciences | 612562 | AB_399853 |
| Goat anti-Mouse IgG1 Cross-Adsorbed Secondary Antibody, Alexa Fluor™ 647 | Thermo Fisher Scientific | A-21240 | AB_2535809 |
| Alexa Fluor™ 488 Phalloidin | Thermo Fisher Scientific | A12379 | n/a |
| Hoechst 33342 | Thermo Fisher Scientific | 62249 | AB_10626776 |
| Image-iT™ FX Signal Enhancer | Thermo Fisher Scientific | I36933 | n/a |
| M.O.M.® (Mouse on Mouse) Blocking Reagent | Vector Laboratories | MKB-2213-1 | AB_2336587 |
| 16% Paraformaldehyde aqueous solution | Electron Microscopy Sciences | 15710 | n/a |
| Bovine serum albumin | Thermo Scientific Chemicals | J65097.22 | n/a |
| Normal goat serum | Lampire Biological Laboratories | 7322500 | n/a |
| Triton X-100 | MP Biomedicals 100 mL | ICN807423 | n/a |
| Fluoro Gel with DABCO | Electron Microscopy Services | 17985-02 | n/a |
| R Project for Statistical Computing | R Core Team | n/a | SCR_001905 |
| BioVoxxel Toolbox | BioVoxxel | n/a | RRID:SCR_015825 |
| Fiji is just ImageJ | Fiji | n/a | RRID:SCR_002285 |
